# Correlative cryo-structured illumination fluorescence microscopy and soft X-ray tomography elucidates reovirus intracellular release pathway

**DOI:** 10.1101/2020.01.13.904623

**Authors:** Ilias Kounatidis, Megan L Stanifer, Michael A. Phillips, Perrine Paul-Gilloteaux, Xavier Helligenstein, Hongchang Wang, Chidinma A. Okolo, Thomas M. Fish, Matthew C. Spink, David I. Stuart, Ilan Davis, Steeve Boulant, Jonathan M. Grimes, Ian M. Dobbie, Maria Harkiolaki

**Affiliations:** Diamond Light Source, Harwell Science and Innovation Campus, Didcot OX11 0DE, UK; Department of Infectious Diseases, Molecular Virology, Heidelberg University Hospital, 69120 Heidelberg, Germany; Micron Advanced Imaging Consortium, Department of Biochemistry, University of Oxford, South Parks Rd, Oxford OX1 3QU, UK; SFR-Santé, INSERM, CNRS, UNIV Nantes, CHU Nantes, Nantes, France; CryoCapCell, 155 bd de l’hôpital, 75013 Paris, France; Division of Structural Biology, The Henry Wellcome Building for Genomic Medicine, Roosevelt Drive, Oxford, OX3 7BN, UK; Department of Infectious Diseases, Virology, Heidelberg University Hospital, 69120 Heidelberg, Germany; Research Group “Cellular polarity and viral infection”, German Cancer Research Center (DKFZ), 69120 Heidelberg, Germany

## Abstract

Imaging of biological matter across resolution scales presents the challenge of preserving the direct and unambiguous correlation of subject features from the macroscopic to the microscopic level. We present here a correlative imaging platform developed specifically for imaging cells in 3D, under cryogenic conditions. Rapid cryo-preservation of biological specimens is the current gold standard in sample preparation for ultrastructural analysis in X-ray imaging. However, cryogenic fluorescence localisation methods are by and large diffraction-limited and fail to deliver matching resolution. We addressed this technological gap by developing an integrated, user-friendly, platform for 3D correlative imaging of cells in cryo-preserved states using super-resolution structured illumination microscopy (SIM) in conjunction with soft X-ray tomography (SXT). The power of this new approach is demonstrated by studying the process of reovirus release from intracellular vesicles during the early stages of infection and identifying novel virus-induced structures.

## Introduction

Cell and tissue imaging methods have progressed rapidly in the past decades and new techniques have provided unprecedented opportunities for accumulating biological insights^1^. Currently, cellular ultrastructure can be successfully imaged to an impressive resolution of typically 2-5nm using scanning or transmission electron microscopy (EM)^2^. However, EM methods suffer from limited penetration depth (up to 0.5μm in transmission EM). For 3D visualisation of thicker samples, such as mammalian cells (which can easily grow beyond 3μm thickness) sectioning and serial reconstruction methods can be employed to overcome this limitation but, they involve chemical treatment of samples or complex cryogenic workflows^3,4^. Cryo-soft X-ray tomography (cryo-SXT) addresses this need for direct mesoscale imaging of cellular ultrastructure in thicker vitrified samples, offering a penetration depth in the order of 10μm to a resolution of a few tens of nanometers^5^. Complementary to both EM and SXT, optical microscopy also provides direct access to information on ultrastructure, organelles and molecular localization within cells^6^. Advances in the field include methods that reach classical optical resolution limits, as defined by the Rayleigh criterion^7^, such as wide-field deconvolution^8^ and confocal laser microscopy^9^. In parallel, new microscopes have been developed that surpass the diffraction limit, collectively termed super-resolution microscopy, embracing a wide range of both near-field and far-field methods^10^.

Further benefits accrue through the combination of imaging methods in correlative schemes. Recently, correlative imaging has contributed significant new insights into a range of cellular processes by providing complementary morphological, structural and chemical information beyond what is achievable through the use of any single technique alone^11^. Most commonly, the process has involved correlating fluorescence microscopy (FM) data with EM^12,13,14^. Correlative light and electron microscopy (CLEM) has been used extensively to date, and recent advances include the addition of super resolution visible-light fluorescence^15–17^. FM data, at cryogenic and non-cryogenic temperatures, have also been correlated with SXT, but in all cases it has either been diffraction-limited^18–22^ or involved a chemical fixation step^23^ with the associated risk for artefact formation^24,25^.

As an alternative to chemical fixation, rapid cryo-freezing is the current gold standard for preserving delicate biological structures or arresting rapid dynamic processes in cells^26,27^ followed by cryo-imaging^28,29^. Vitrification was first identified as a promising technique for biological sample preservation for ultrastructural imaging forty years ago^30^ and since then has developed into the method of choice for sample preparation for both X-ray imaging^22^ and electron tomography^27^ and has also been used for super resolution fluorescence imaging^31^. Vitrified samples suffer no fixation artefacts and cellular structures are preserved faithfully in their pre-vitrification state^29^. Fortuitously, fluorescence imaging at cryogenic temperatures is characterised by enhanced photo-stability of dyes and sharper emission spectra as compared to room temperature imaging leading to data with better signal to noise ratios^14,31–33^. However, numerical aperture (NA) in cryo-FM systems is limited because there are currently neither suitable immersion fluids nor robust dipping objectives. As a result, cryo-fluorescence imaging is done using air objectives with a maximum NA of circa 0.9, limiting the achievable optical resolution. Moreover, it remains technically challenging to prevent ice crystal formation and sample damage while imaging under cryogenic conditions. Samples have to be kept below the glass transition temperature of circa 130°K (−140°C), whilst mounted close to an objective that is kept at room temperature and while at the same time being illuminated with high intensity light for excitation purposes. Given these challenges, 3D-structure illumination microscopy^34,35^ is the natural choice for a super resolution method for cryogenically preserved biological samples as it can deliver fast 3D super-resolution imaging with relatively low exposures to laser light and wide fields of illumination.

Here, we present a novel correlative imaging scheme for biological samples at cryogenic temperatures that uses a purpose-built 3D-cryoSIM microscope^34,36^ and a synchrotron soft X-ray microscope capable of tomographic imaging. These are available as an accessible and user-friendly imaging platform at beamline B24^5^ at the national UK synchrotron, Diamond Light Source (DLS; http://www.diamond.ac.uk). The techniques implemented are highly complementary and deliver comprehensive views of both cellular ultrastructure and molecular organization over extended cellular volumes in minimally perturbed cell populations. We have applied this novel combination of imaging capabilities towards the investigation of the early events of reovirus infection in mammalian cells and in particular the mechanism of viral penetration of endocytic vesicles.

## Results/Discussion

To address the current limitations of cryo-imaging relatively thick biological samples at a near-physiological state to tens of nanometres resolution we have developed a correlative imaging platform for data collection on cryo-preserved vitrified samples (primarily cell populations, adherent or in suspension) deposited on gold 3mm EM grids (see **Methods**; **Supplementary Figure 1**). Grids are first mapped using conventional brightfield cryo-fluorescence imaging where the areas of interest are identified before 3D data is collected, first using our newly developed cryo-3D-SIM system and then at a soft X-ray full-field transmission microscope. Data are then processed and correlated to produce 3D imaging volumes that contain both chemical localisation and ultrastructural organisation information enabling the meaningful and unambiguous interpretation of biological features.

### 3D cryo-Structured Illumination Microscope (cryoSIM)

Imaging data can convey information beyond the diffraction limit of a microscopy system via super resolution methods, a field that has expanded rapidly over the last 20 years^29^. This has revolutionised biological imaging leading to a number of outstanding bespoke instruments as well as off-the-shelf commercial microscopes^37–41^. Structured illumination microscopy, in specific, is a 3D FM super resolution method^34^ that not only delivers an 8-fold increase in volumetric resolution as compared to the diffraction-limit but, also has distinct advantages over other super-resolution techniques: (a) it is easily applied to multi-channel imaging using a range of common dyes or fluorescent proteins, opening up a wide range of biology for study; (b) it can image relatively thick samples, reducing the need for sample sectioning prior to imaging; (c) it uses relatively low light doses (10-100 W/cm^2^ laser power) and short image acquisition times (20-100ms per single exposure), minimising sample heating and reducing the chance of ice crystal formation which would otherwise lead to sample damage and image quality degradation^35^ and, (d) it has extremely good out of focus light suppression, leading to high contrast images even in samples with thicknesses of 10μm or more^36^.

Given the current requirement to extend the application of super-resolution methods to the study of cryogenically preserved samples for the purposes of correlative imaging, we present here a novel 3D-SIM setup^39,40^ with an open optical design that overcomes many of the technical difficulties of 3D imaging fluorescence at super-resolution under cryogenic conditions. The conceptual point of reference for our design was the OMX platform^41^ (designed and built by Prof. John Sedat and his team^34^).

This custom-designed cryoSIM is built at beamline B24 around a commercial cryo-stage (CMS196M, Linkam Scientific) with a long working distance air objective (100X, 0.9NA, 2mm working distance, Nikon) and delivers imaging at a maximum lateral resolution of circa 360nm for green fluorescence detection (see **Methods**, **Figure 1a-c** and **Figure 1e-m** for optical performance with nanobeads and cells, respectively). By using a relatively high NA long-working-distance air objective, we are able to maintain samples at cryogenic conditions (71K) without dipping the objective in cryogen. The system, currently, has four illumination wavelengths (405, 488, 561 and 642nm) from individual direct-diode and diode-pumped solid state lasers, which could, if necessary be modified to include any other separable channels in the visible wavelength range. A nematic liquid crystal spatial light modulator is used in phase modulation mode to produce structured illimitation (SI) stripe patterns, and a liquid crystal based polarisation rotator (Meadowlark Optics) allows us to optimise SI patterns independently for each stripe orientation and excitation wavelength, eliminating the need to optimise for one to the detriment of all others (see **Methods**). The system’s optical features are incorporated under the label ‘cryoSIM’ in the available-online Spekcheck tool^42^ where the efficiency and performance of different fluorophores can be assessed *in silico* with respect to features in our setup, thereby enabling the intelligent design of fluorescence-dependent imaging experiments (**Figure 1d**). 3D-SIM requires 30 times as many images as widefield fluorescence, 15 images per Z-plane, and twice as many Z-planes to satisfy Nyquist-Shannon sampling^7^. At the cryoSIM, each Z slice is recorded 15 times (five phases for each of three angle in the SI pattern) at 125nm increments along the Z axis as 512×512pixel images with 125nm per pixel and reconstructed to 1024×1024pixel image stacks with voxel size of 62.5×62.5×125nm (see **Methods**). Point spread functions for reconstruction purposes at cryogenic temperatures are generated before data collection using 175nm *PS-Speck* nanobeads (ThermoFischer) at emission wavelengths of 440, 525, 605 and 647nm.

**Figure 1;.**
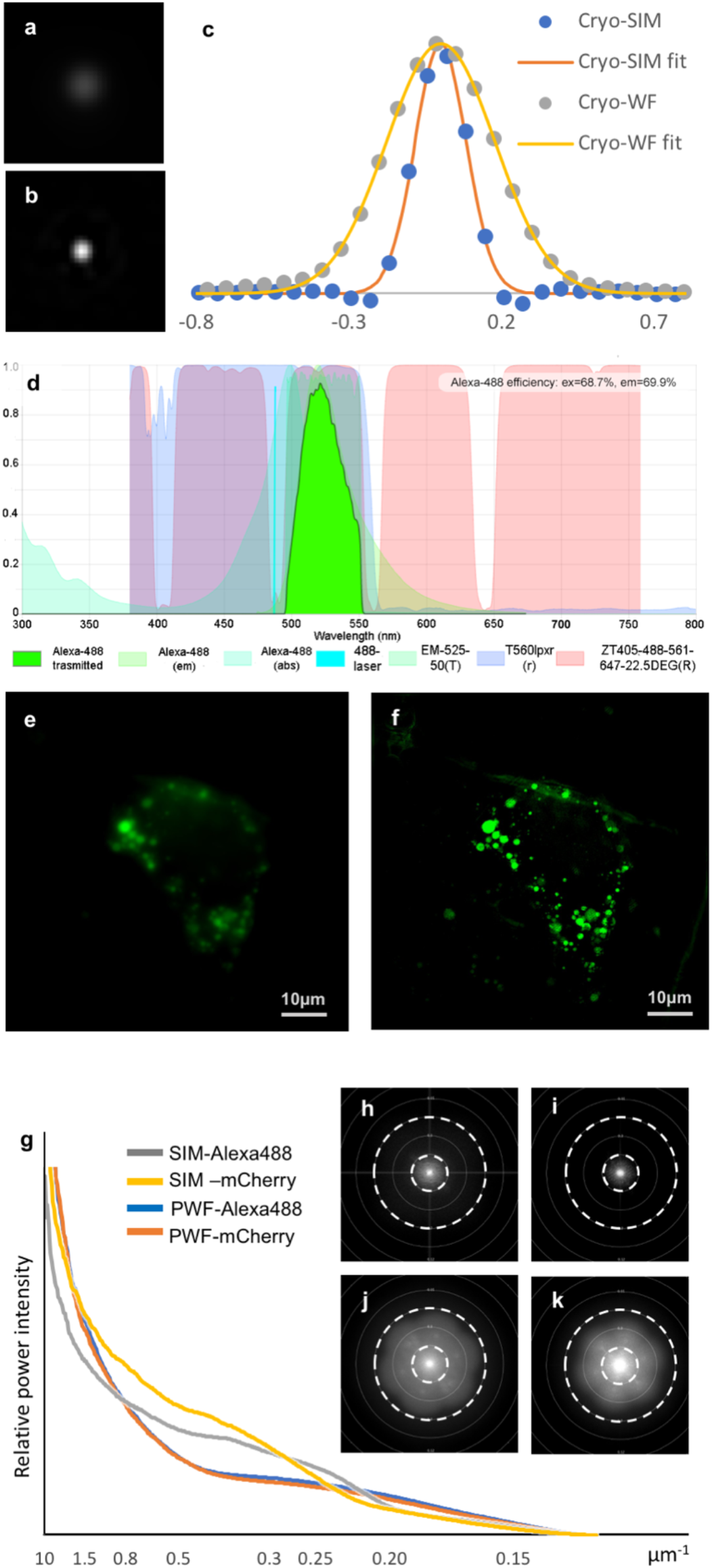
CryoSIM optical performance. **a**, Widefield and **b**, SIM lateral point spread of a 175nm diameter, 505/515nm wavelength microsphere (PS-SpeckTM ThermoFischer) collected on the cryo-SIM at 71K using 488nm light for excitation and a 520/25nm filter (Semrock) for emission. **c**, Plot of lateral point spread (single line profile) of **a**, and **b**, and their respective gaussian fits showing clear resolution enhancement. **d**, SPEKcheck^42^ display of the cryo-SIM for a representative fluorophore (Alexa488) excited in the system with an efficiency of 68.7% and a resulting collection of 69.9% of the total emitted light. Laser light is shown as a bright blue spike, excitation and emission are shown in shades of light green and the total transmitted light is shown in bright green. The system dichroic mirrors and filters are also represented (system dichroics denoted as ZT, detector splitter dichroic as T560 and the emission filter as EM). Analysis of representative cryo-SIM data from a single mammalian U2O9 cell containing structures tagged with Alexa488 fluorophore: **e,** Z axis projection of the raw SIM data processed to produce a pseudo-widefield image stack; **f,** Z axis projection of the same SIM data fully reconstructed showing the resulting resolution enhancement. **g**, Intensity analysis of plots of **h-k** (flattened and sampled radially) showing the relative increase in information content after reconstruction. **h** and **i**, Reciprocal space resolution plots of the raw pseudo-widefield data in the cell shown in **e** (Alexa488 fluorescence and mCherry fluorescence (associated with a tagged reporter molecule) respectively). **j** and **k**, Same data as **h** and **I** respectively after SIM reconstruction. Concentric dashed circles for **h**-**k** denote resolution boundaries of 600nm (small circle) and 200nm ((large circle). All image analysis was done using Fiji^43^ and the SIMcheck^44^ Fiji plugin.

### Cryo Soft X-ray Tomography (cryo-SXT)

The full field transmission X-ray microscope (TXM; Zeiss UltraXRM-S220C) is a synchrotron endstation installed at beamline B24 (DLS, zone 4), and delivers imaging by absorption contrast. The TXM allows imaging within a defined spectral region known as the ‘water window’, which lies between the k absorption edges of carbon at 284eV and oxygen at 543eV. Within this region, carbon-rich biological structures absorb X-rays more than the surrounding oxygen-rich medium and the resulting natural contrast allows the delineation of cellular features in recorded projections. Cellular membranes provide particularly strong contrast, making membrane bounded organelles especially easy to identify within relatively thick ultrastructural 3D tomograms of whole cells

The TXM is illuminated by synchrotron radiation supplied by a bending magnet, focused by a toroidal mirror and conditioned with a plane grating monochromator and exit slit module that can deliver highly monochromatic beam at 500eV (**Figure 2a–2c**). This beam forms a secondary light source which is delivered to the microscope and focused by a glass capillary condenser lens onto the sample. A zone plate objective focuses the resulting projections onto a highly sensitive photon detector (Pixis1024B CCD, Princeton Instruments). Samples on standard cryo-EM grids are placed into the X-ray microscope using a transfer chamber that facilitates transition from liquid nitrogen storage under atmospheric pressure to active cooling via conduction in high vacuum (10^-6^ to 10^-8^mbar, essential as soft X-rays have poor penetration at atmospheric pressure). Because of the limited divergence of synchrotron bending magnet sources, the size of the focused X-ray beam at the sample position is approximately 1.2μm. To illuminate the maximum field of view (FOV), the condenser follows a Lissajous curve and, to match the 40nm zone plate objective fills a 16×16μm^2^ area at the sample plane (10×10μm^2^ for the 25nm objective) (**Figure 2d**). Samples at the imaging position are close to both the capillary condenser and the zone plate objective, circa 6mm and 5mm from each^42^ (**Figure 2e**), limiting specimen tilt to a maximum of ±70°, which leads to missing wedge artefacts. This may be mitigated in part by tilting around two orthogonal axes^45^. The zone plate objectives of the system are designed to focus soft X-rays; they are made of a series of concentric metal rings of radially decreasing width that are installed approximately 5mm after the sample (beam divergence of 1.7mrad; zone plate diameter of 150μm. The resolution, *δ*, of the microscope depends on the width of the outer most zone width *(δ* = 1.22 Δ*r_n_* for coherent light) and there is a choice between a 25nm and a 40nm minimum width for absorption contrast imaging (depth of focus of circa 1 and 3.5μm respectively) (**Figure 2f-g**). The lower resolution delivered by the 40nm zone plates is accompanied by an increased FOV and depth of focus making it well suited for studies aimed at collecting the maximum amount of ultracellular structure, such as for example the study of organelle shuttling and restructuring. The TXM delivers magnification up to 1,300x with a square FOV of view of between 10 or 16μm across and records 1024×1024 pixel projections at either 10nm or 16nm per pixel, depending on the objective in use (25nm and 40nm respectively). In addition, we have basic in-line visible light microscopy capacity with broad-spectrum LED illumination (widefield and fluorescence) through a 20x objective located within the sample chamber to allow mapping and registration of regions of interest (ROIs) previously identified by conventional or super resolution light microscopies (see below). Samples are vitrified before exposure to X-ray radiation to immobilise them and preserve them against radiation and heat damage. Gold nanospheres, 100nm-250nm in diameter, are traditionally included in sample preparation just before vitrification to provide highly absorbent points of reference (fiducials) for image alignment necessary in data reconstruction. 3D imaging data is collected as a series of projections at defined regular angular steps generating of a tilt series up to 140° in total with projections collected typically at 0.5-1s exposures. The images are subsequently processed into tomograms using IMOD^46^ through a fully automated, *in silico* pipeline that runs concurrently with data collection (see **Methods**). To capture cellular content extending beyond a single FOV, multiple tomograms can be collected at adjacent overlapping areas. For example, when vesicles and substructures of interest need to be captured along with the overall cellular landscape in which they reside.

**Figure 2;.**
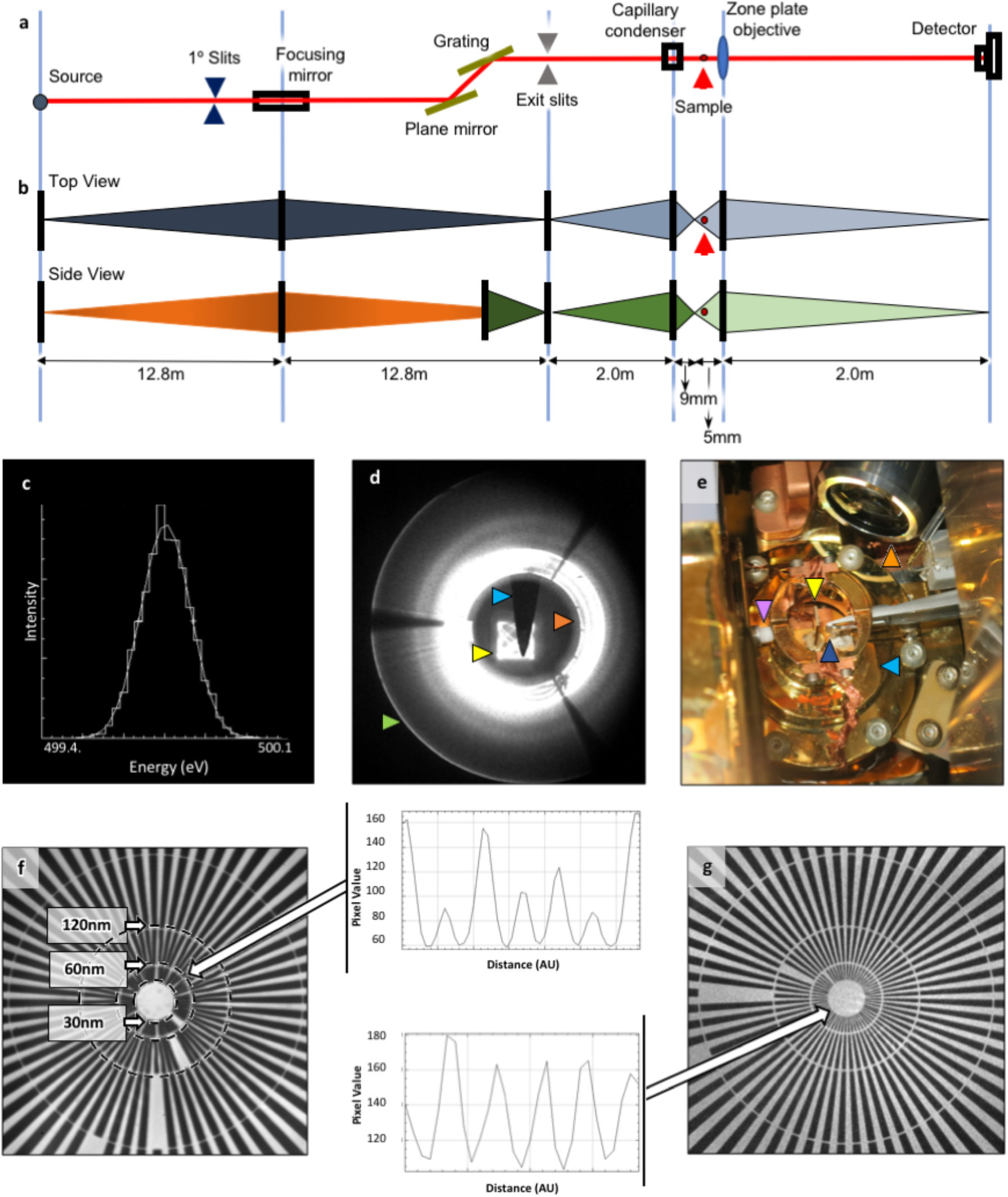
TXM characteristics. **a**, Schematic representation of the X-ray optical path at beamline B24. **b**, Beam divergence profile and monochromacy, top and side view. The bending magnetic source light is focused by horizontal deflection of 1.1° with 1:1 magnification by a toroid focusing mirror located 12.8m downstream. A plane mirror-grating monochromator directs the real image of the source light, now monochromatic, a total of 25.6m from the bending magnet it originated from. This beam is then focused by a capillary condenser to the sample and a zone objective delivers the magnified image to the detector. **c**, Energy profile in the beam that illuminates the sample. **d**, Imaging beam profile recorded at a scintillator 0.5m downstream from sample position; the focused beam is used to fill a central square FOV (yellow arrow) with a Lissajous light pattern the result of a two-motor condenser sweep, with light reflected from the condenser at the maximum angle (green arrow) and the inner surface of the condenser (red arrow). An alignment Tungsten pin (blue arrow) can be seen at the sample position in the middle of the illuminated area. The image was acquired on an alignment camera via a scintillator positioned 0.5m downstream from sample position. **e,** View of the sample environment in the TXM showing the condenser (purple arrow), the objective (dark blue arrow), sample holder (yellow arrow), the 20X visible-light objective (orange arrow) and cryo-shield (light blue arrow; needed to minimize thermal drift between the 71K ‘cold’ sample and the optics which are all maintained at 21°C; sample chamber maintained at high vacuum >10^-6^Torr. **f** and **g** X-ray projection of a 30nm (minimum spacing) Siemens star (Xradia Inc, now Zeiss) at beamline B24 imaged with 40nm and 25nm objectives respectively; resolution is measured by observable dips in absorption at >2sigma using a line profile across several spokes at the inner edge of each concentric zone. Arrows emanating from the associated graphs show the resolution edge the line profile was collected at. Dotted circles denote the edge of each zone and the spacing therein. Line profiles were generated with Fiji^43^ (vertical axis shows pixel intensity values; horizontal axis shows representative spacing in arbitrary units (AU), not calibrated against the real distance which is denoted by the corresponding dotted circle). **f** and **g,** show that the TXM delivers resolution beyond 60nm and 30nm resolution with the 40nm and 25nm zone plate objectives respectively.

It is important to note that, deciding on a data collection strategy before acquisition is vital. Parameters such as sample thickness and susceptibility to radiation damage (heating due to expose to X-rays) need to be balanced against optimal single frame exposure time and sampling step for the resolution required while sampling of multiple overlapping FOVs is entirely dependent on how robust the sample is to extended exposures. Moreover, background noise is accumulated in the raw data images due to sample holder materials such as thin carbon support film, soluble intracellular content or surrounding leftover cell media in combination with sample or beam mechanical instabilities during data collection can reduce contrast in the data and interfere with the delivery of the best achievable resolution (**Figure 2f** and **2g; Supplementary Figure 2**)

### Experimental workflow of correlative cryo-SXT and cryo-SIM

Cryopreservation lies at the heart of the workflow as it allows not only the immobilisation and preservation of native cellular structures but also confers resistance to prolonged exposure to intense light during imaging. Cells to be imaged are typically grown as adherent monolayers, multilayers or in suspension. Adherent cells are cultured on carbon films on gold transmission electron microscopy (TEM) grids^47^ ideally with positional markers (finder grids). Fluorescent markers can be endogenously expressed or added to the culture media for uptake into the cells. Attachment, growth, confluency and distribution of live cells in the media can be established via any number of conventional light and fluorescence microscopy methods prior to addition of fiducial markers (usually gold nanobeads), blotting (to remove excess media) and vitrification via plunge freezing in liquid nitrogen-cooled liquid ethane. Plunge freezing^48^ is the current method of choice for sample preservation for cryo-SXT, with high pressure freezing being required for samples thicker than 10μm^49^. Post-vitrification, grids are once again imaged in both brightfield and fluorescence and mapped at the beamline on a cryostage-equipped light microscope with fluorescence detection capabilities, to examine cell morphology and to confirm absence of non-vitreous contaminants, and mapped at low resolution (Linkam Scientific CMS196M; Zeiss Axio Imager 2; DIC 50x, 0.55NA). At this point, the selected samples are ideally intact, with thin vitreous ice around them and ROIs have been identified. Grids can now either be used as they are or clipped in autogrid holders (ThermoFisher) for greater stability. The cryo-SIM is thereafter the first stop for high-resolution 3D imaging, followed by cryo-SXT on the same ROIs. As exposure to soft X-rays leads to atomic bond damage and irreversibly degrading fluorophores and hence X-ray imaging always follows fluorescence imaging.

Samples are placed in standard Linkam cryoholders at the cryoSIM which is initially used to generate a brightfield transmission 2D mosaic of the grid holder structure and ROIs are annotated on this mosaic as data collection takes places. 3D-SIM data are collected at specified Z extends (4-12μm depending on cell thickness at Z-intervals of 0.125μm). To achieve optimal resolution enhancement, the cryoSIM captures three angles and five phases of stripes, leading to a total of 15 images per Z slice. Each exposure depends on overall fluorophore intensity, but commonly we expose for tens of milliseconds using laser light at 10-100mW/cm^2^ illumination power. A typical data set is collected within an average time of 3-5min capturing a FOV of 64×64μm in all Z slices required. We have not observed any thawing of vitreous samples during data collection under the above-mentioned standard conditions and samples that have been used in cryoSIM imaging do not appear structurally different from other samples at the TXM. After SIM imaging samples could also be used for cryo-electron tomography through a focused ion beam milling step^50^, however, we have yet to fully explore this potential.

Following cryoSIM data collection, grids are recovered from the Linkam holders and stored until they can be loaded into TXM holders (either conventional Zeiss cryo-holders, or autogrid™ TXM holders the design for which was a kind gift by the Mistral beamline at the Spanish synchrotron, ALBA [https://intranet.cells.es/Beamlines/XM]). Once at the TXM, grids are examined and mapped with visible-light to produce a new mosaic at the sample orientation (moving from one holder to the other results in random sample rotation in each microscope). The two mosaics can be aligned with a semi-automated plugin at the beamline and the established ROIs from SIM data collection are recovered to allow X-ray data collection in the same regions. The samples are further evaluated for radiation damage and possible freeze-thawing (semi-crystalline ice areas can be readily identified in X-ray projections) and a data collection strategy is devised based on trial exposures in areas away from established ROIs in conjunction with the imaging requirements of each project. 2D X-ray mosaics are acquired at all ROIs covering single grid squares (imaging of 7×7 adjacent FOV at 16×16μm will image a whole grid square in a standard EM grid) and these are used to decide on data collection areas. 3D X-ray data is collected as a tilt series as described previously. They are relayed in real-time to an automated pipeline using IMOD Batchruntomo^51^ and, provided a sample has good contrast and fiducials (ideally >3 and well dispersed in the FOV), the data are reconstructed to tomograms using IMOD’s weighted back projection, serial iterative reconstruction and batch processing options. The resulting tomograms can then be used to further refine data collection strategy. Processed data from both high-end microscopes can then be used in eC-CLEM^52^ for the *in-silico* 3D correlation of associated features, 3D overlay and further analyses (see **Methods; Figure 3; Supplementary Figures 3 and 4**).

**Figure 3;.**
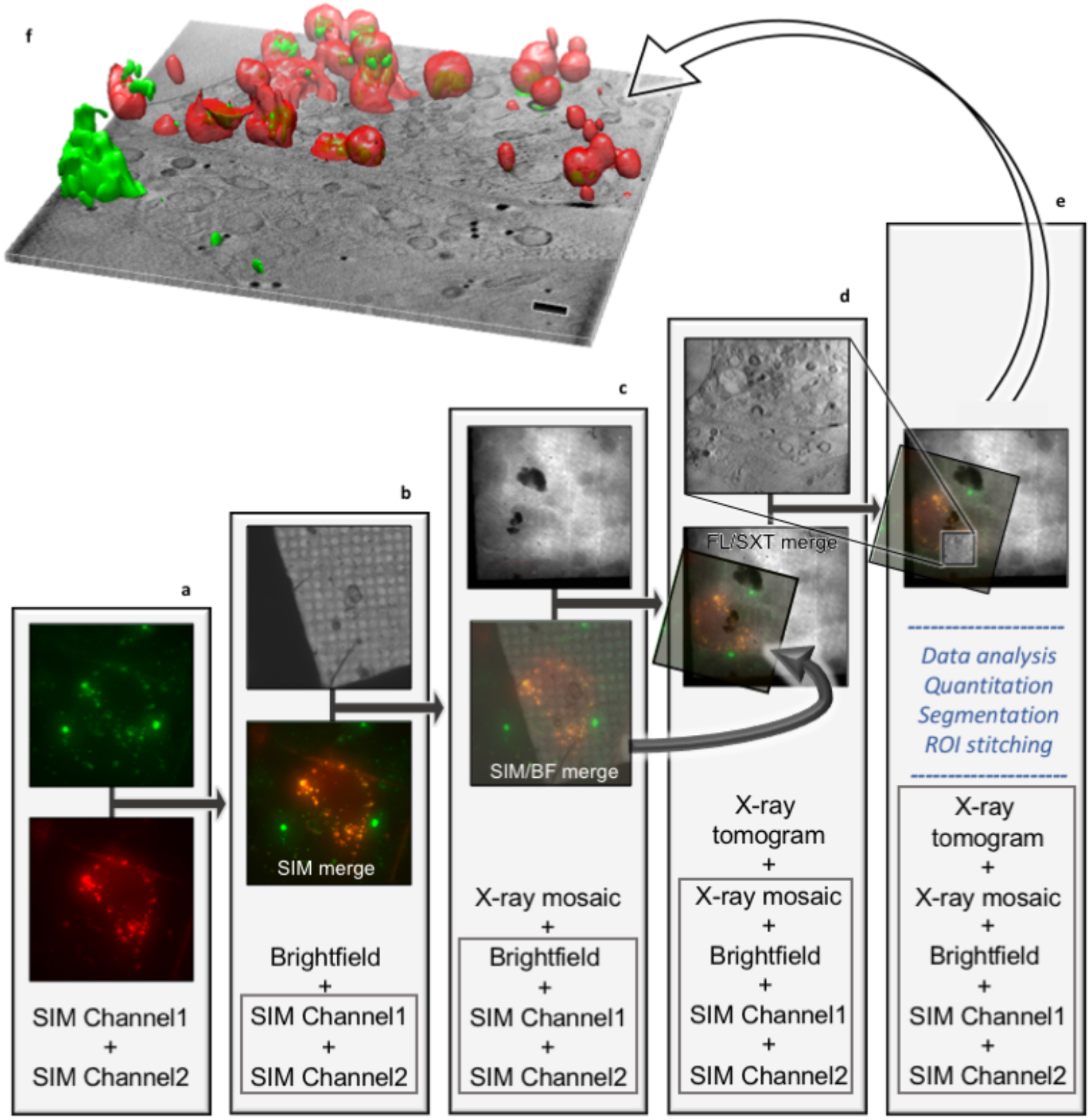
Stepwise protocol for in silico correlation of data collected at beamline B24. **a**, Correlation of fluorescence signals on the same ROI, **b,** superposition of all fluorescence signals (bottom) with corresponding brightfield data (top), **c**, positioning of combined image stacks from previous step (SIM/FM merge; bottom of column) in 2D X-ray mosaic (top), **d**, 3D superposition of previous (bottom of column) with X-ray tomogram at ROI (top; lateral resolution of imaging beyond 60nm; measured resolution at the B24 transmission X-ray microscope with the 40nm objective). **e**, The combined 3D stacks can now be further analysed with image analyses packages such as Fiji^43^ or segmentation algorithms such as SurVoS^53^ to extract statistical data on cellular structures or further combined with tomograms from adjacent ROIs to increase the 3D space imaged or segmented. **f**, The resulting multi-channel stack has all imaging data correlated in 3D space (a single slice from the middle of the tomograms is shown with the correlated 3D fluorescence volumes (red for red fluorescent intracellular vesicles that contain molecules of interest that fluoresce green; the scale bar is 1μm). Superposition in steps **a** and **b** were done with Chromagnon^54^ and **c**, **b** and **e** with eC-CLEM^52^; visualisation and rendering were done with Fiji^43^ and Chimera^55^.

### Proof of concept application of 3D cryo-correlation workflow: Elucidating early events in reovirus infection

Viruses are infectious particles, which must enter a host cell to propagate. Therefore, crucial to any viral infection is the passage through the physical barrier of host membranes. For enveloped viruses, the mechanisms of virion delivery are relatively well understood and rely on fusion of the lipid bilayer of the viral envelope with the lipid bilayer of the host^56^. Reovirus, however, as a non-enveloped icosahedral virus, does not have membrane fusing capacity and how its core reaches the cytoplasm to initiate replication has remained largely elusive. Reoviruses have a segmented double-stranded RNA genome, which during viral infection is retained within the closed core particle, preventing the triggering of cellular innate immune responses. Viral core particles have both the RNA polymerase and the capping enzyme required for transcription of viral mRNA and viral replication. The core particle of reovirus is cloaked by the outer capsid, which is composed of proteins σ3 and μ1, plus the σ1 spike protein at the icosahedral 5-fold axes. σ1 and σ3 are involved in receptor attachment whilst μ1 plays a role in viral entry^57^. Reoviruses have been shown to enter cells via clathrin-mediated endocytosis and are subsequently trafficked through the endosomal pathway from the early Rab4,5 or 11 positive endosomes to the late Rab7 or 9 positive endosomes^58,59^. The escape of reovirus core particles from these endosomes to the cytoplasm has been linked to low-pH-dependent proteolytic degradation of the viral outer capsid protein σ3^60^. There is *in vitro* evidence that proteolytic removal of σ3 (likely to occur in endosomes due to gradual increase in acidity and digestion by low-pH dependent-cathepsins^61^), results in the release of a N-terminally-myristoylated μ1 peptide, which can result in pore formation^62^. There are still, however, important unanswered questions regarding the mechanisms driving the early stages of reovirus infection, such as: (a) how soon after entry are replication-competent core particles released into the cytoplasm, (b) are they released at specific locations within the cytosol and (c) is the ultrastructure of endosome vesicles affected by viruses during their entry, trafficking and release? Our current lack of understanding of the details of core particle intracellular release is due in large part to the absence of technological solutions to visualize, at sufficient resolution, the cellular landscape during infection and document its remodelling in response to viral infection.

To better understand reovirus escape from endocytic vesicles, we have used the human bone osteosarcoma U2O9 cell line expressing mCherry Galectin-3 (U209-Gal3; see **Methods** for experimental details). Gal3 binds to extracellular carbohydrates and within a normal cell, is distributed throughout the cytoplasm as it is produced and is then trafficked to the cell surface. However, upon endosomal membrane disruption, extracellular carbohydrates that were internalized through endocytosis, become accessible to Gal3 as it floods in along with other cytosolic proteins, and binds to carbohydrates within the disrupted endosomes. Hence, Gal3 serves as a useful indicator of endosomal membrane disruption^63^.

To map the time course of reovirus localisation, U209-Gal3 cells were transfected with Rab7-GFP (late endosomal marker) and subsequently infected with fluorescently labelled T3D reovirus. Live-cell spinning-disc confocal microscopy was used to follow virus entry, the association of the fluorescent virus with the late Rab7-marked endosome and the appearance of localised Gal3 upon endosomal rupture process. Our data show that, within 1 hour post infection (PI) more than 60% of virus particles are associated with Rab7 positive endosomes (**Figure 4a-b**). Importantly, by 2 PI 60% of Gal3 positive vesicles are associated with virus particles (**Figure 4c**). U209-Gal3 cells were then seeded on TEM gold grids following infection with high titres of Alexa488-labelled T3D reovirus (see **Methods**). Grids with infected cells were lightly blotted to remove excess media and plunge frozen into liquid nitrogen-cooled liquid ethane at distinct time points post exposure to the virus (T=0, 1,2 and 4h). All grids were first mapped through conventional cryo-microscopy in accordance with the workflow previously discussed (**Supplementary Figure 5**) and ROIs were 3D imaged first using the cryoSIM (see **Methods**). The resulting super resolution imaging data extended the confocal study by capturing small Gal3-positive vesicles that are also positive for reovirus by 1h PI (**Figure 4d-g; Supplementary Figures 6 and 7**), and these structures are seen to further increase in number, size and signal intensity by 2h PI, gradually forming large vesicle-like perinuclear structures (up to 3μm diameter) by 4h PI (**Figure 4h-k; Supplementary Figures 6 and 7**). It is therefore evident that, by 1h PI, endocytosed reovirus particles have started disrupting endosomal membranes sufficiently to allow an influx of Gal3 and the affected endosomes continue to progress to the late stages of development. Therefore, we now have a clear timeline for viral escape (1-2h PI) and clear virus transport through conventional endosomal transfer.

**Figure 4.**
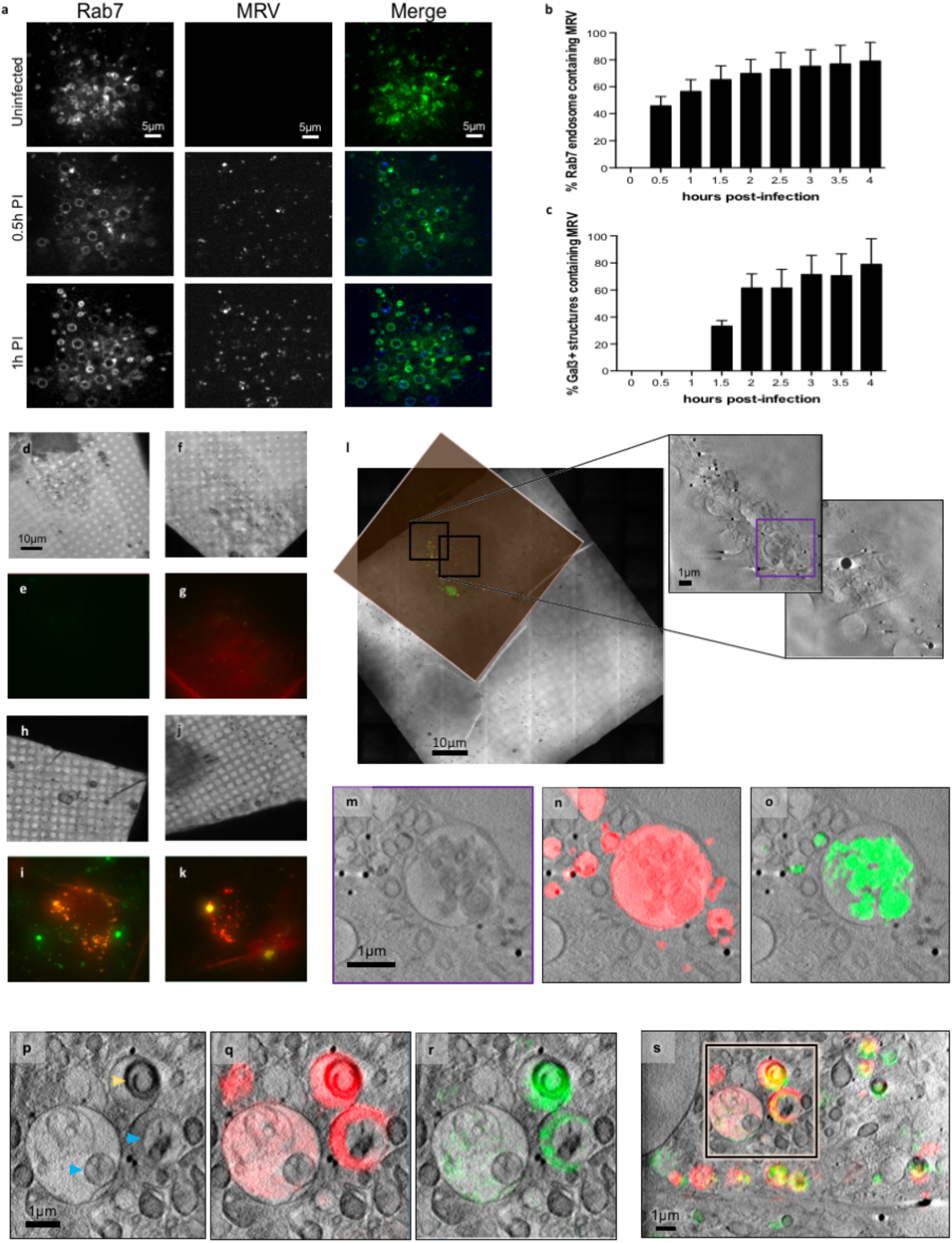
Tracking reovirus endosomal trafficking and escape. **a,** Confocal images of U209 cells expressing late-endosome marker Rab7-eGFP infected with Alexa647-labelled reovirus at indicated times PI. **b**, Mammalian reovirus (MRV) presence in late endosomes up to 4h PI and **c**, numbers of Rab7-positive endosomes containing MRV for the same period (error bars indicate standard deviation of n=8 in **b** and n=9 in **c**). **d-k**, CryoSIM images of MRV labelled with Alexa488 in mCherry-Gal3 expressing cells showing the concentrations of Gal3 signal in distinct vesicles in red; brightfield and SIM slices respectively at: time 0 mock-uninfected (**d** and **e**); 1h PI (**f** and **g)**; 2h PI (**h** and **I**); 4h PI (**j** and **k**). **l**, Superposition of SIM processed data of a 4h PI sample on the 2D X-ray mosaic of the same area and expansion of adjacent FOVs where two adjacent X-ray tomograms were collected and stitched. **m**, Expanded region of interest (highlighted purple in l). **n** and **o**, same as **n**, with Gal3 signal in red and reovirus signal in green respectively. **p**, **q** and **r**, ROI from 2h PI: single Z-axis slice from the corresponding tomogram with Gal3 (red) and MRV (green) localisation. Multi-vesicular bodies (MVPs) are clearly identifiable with distinct sub-compartments where Gal3 is concentrated (presumably the result of virus-carrying vesicles fusing with virus free late endosomes). Novel virus-induced carbon-rich substructures are marked by yellow arrows and vesicle areas within MVBs that remain impermeable to Gal3 marked by blue arrows. **s**, Overview of the relevant 2h PI region with both fluorescent signals overlaid.

Having established the timeline of viral disruption of endosomes using cryo-SIM (corroborated by our previous live cell confocal microscopy under similar conditions), we needed a high-resolution view of the landscape within infected cells that would unambiguously give us the 3D characteristics of the carrier organelles involved in virus trafficking; their structure, possible virus-induced damage and associated relative distribution. To achieve this goal, grids carrying vitrified infected cells, with ROIs already imaged at the cryoSIM were first mapped via the inline visible light brightfield and fluorescence microscope at the TXM followed by imaging via SXT (see **Methods**; **Supplementary Figure 8**). For this project, we chose to image using the 40nm objective of the microscope taking advantage of the extended FOV it affords us in order to collect large volumes of data and provide suprastructure context for the vesicles. The alternative of higher resolution imaging would have allowed us to potentially identify individual virus particles in the cytoplasm but over much reduced areas within cells (this high resolution aspect of the project is currently ongoing). SXT allowed us to unambiguously delineate vesicle structures across the cytoplasm including areas very close to the nucleus which would be largely inaccessible to other established nanometer-resolution techniques such as electron tomography (**Figure 4m-s; Supplementary Figure 9; Supplementary Movie 1**). Using SXT we observed: (1) populations of simple vesicles carrying viral load at all time points examined and (2) some vesicles progressively growing larger in size and changing to multi-vesicular bodies (MVBs) localised away from the cell periphery and closer to the nucleus. MVBs were visible from 2 hours PI (**Figure 4p-s**), onwards and continued to grow in size and complexity throughout the 4h period after exposure to reovirus. We also observed (3) a novel subpopulation of vesicles with polarised carbon content distribution forming distinct ‘horseshoe’ structures **(Figure 4p, Supplementary Figure10)**. These structures were only observed in infected cells 2h PI and were not observed at any point in the cytoplasm of control cells (**Supplementary Figure 9**), suggesting that they are formed in response to viral presence. It is important to note here that, all cells were exposed to high titres of a virus reputed for its high infectivity and, although this is not necessarily dissimilar to the virus prevalence during an established infection physiologically (an infected cell can deliver hundreds of virions in its immediate environment) we cannot exclude the possibility that some of our observations might be the result of this ‘overloading’. Hence, the above mentioned ‘horse-shoe’ structures could be the result of the accumulation of viral waste products because of the large numbers of particles intracellularly or it could represent a pathway for viral extracellular release and can be studied further in the future.

Once all data were collected, we used eC-CLEM^51^ to align in 3D all associated imaging which allowed us to expand the information content available (**Supplementary Figures 11-14**). Through this complimentary approach, we confirmed that reovirus localised in smaller vesicles (presumably endosomes) from 1h PI which develop into larger vesicles by 2h. These observations suggest that initial engulfment is followed by smaller vesicles merging to form progressively larger vesicles while their outer membrane carbon absorption remains relatively uniform at different time points. From 1 h onwards we also confirmed the presence of localised Gal3 within virus-bearing vesicles suggesting that their membranes have been modified enough (presumably through pore formation) to allow influx of cytoplasmic proteins and possibly other small cellular constituents. Crucially, the implication of this is that core particles are able to escape to the cytoplasm at this early PI. We can now confirm that these events do not disrupt the overall ultrastructure of virus-laden vesicles since none of the Gal3 and reovirus positive vesicles appear misshapen or deflated and have no ruptures of their membranes which are contiguous across the circumference of each vesicle. Assuming that virus-induced pores shuttle viral cores (a sizeable 60nm or more) across the endosomal lipid bilayer, the observation that the integrity of the affected endosomes is not structurally compromised is suggestive of selective shuttling of core particles with pores closing or reducing in size when not in use to ensure endosomes remain apparently undamaged. This would further aid in shielding the virus from detection during the early stages of infection. By 2h PI virus-positive endosomes are becoming larger, more complex and multi-vesicular in structure and are found closer to the nucleus, indicating that they are trafficked for either recycling within or expulsion from the host cell. It is noteworthy that, using our correlative approach, the distinct dome like ‘horse-shoe’ structures observed with SXT within single vesicles in the cytoplasm by 2h PI (**Figure 4p; Supplementary Figure 10**) can now be unambiguously identified, via the cryo-SIM data correlation, as virus associated, suggesting that they are vesicle-sequestered structures likely enriched with the remnants of reovirus outer capsids.

It is important to note that our resulting correlative data is rich in content (**Supplementary movies 2** and **3**) that could not have easily been acquired by other microscopy methods without sectioning or other potentially artefact-inducing sample processing. We have demonstrated here that this correlative approach is powerful and likely to be generally applicable. While each of the imaging modalities described here contributes independently to our understanding of the system under study, once combined within the unique correlation scheme presented here, they allow us to characterise and further understand medically-relevant biological process such as the early events of viral infections **(Supplementary Figure 15)**. Future work will concentrate on further analysis of biomarkers on the endosomes under study, tracking of lysosome merging and the full characterisation of the novel single vesicles observed at 4h PI as well as their origin and possible role in infection or pathogen clearance.

### Concluding remarks

Cryo-SXT is currently the only method that allows the visualisation of cellular substructure through thick samples such as the perinuclear area of fully hydrated cells without the need of fixation, staining or sectioning and the potential associated artefacts, especially likely for membrane structures such as endocytic vesicles. Using this method, we were able to collect clear 3D representations of carbon-rich cellular structures within a reovirus-infected cell population. To complement the information content of our tomograms, we selected 3D-SIM as the super resolution imaging technique of choice as it has distinct advantages for cryo-imaging of biological samples suitable for cryo-SXT. It requires relatively low illumination levels while still giving up to twice the resolution of conventional widefield or confocal fluorescence imaging. It is also able to image 3D samples over a relatively large depth and has the lowest total light dose among the super resolution techniques. SIM imaging can be performed straightforwardly using standard dyes, or fluorescent fusion proteins, and it is simple to image a single sample using multiple fluorescent channels. Using cryo 3D-SIM we were able to unambiguously identify which endosomes within an infected cell carry virus, and when the viral cores likely escape these vesicles. Interestingly, the observation that these virus-laden endosomes maintain structural integrity gives credence to a controlled pore-facilitated penetration mechanism rather than escape via rupture of endosomal membranes. The distinct ‘horse-shoe’ carbon-rich structures observed in vesicles only after virus infection have been shown to be virus-enriched and are either structures likely enriched with the remnants of outer capsid proteins left behind after the release of reovirus core particles in the cytoplasm or a way of whole virus shuttling that has not been observed previously.

This study demonstrates the power of correlative imaging approaches in revealing structural and dynamic properties of complex cellular processes down to a resolution of tens of nanometres. Cryo 3D SIM-SXT correlation brings together two powerful volume imaging techniques that will become invaluable for investigations in a wide range of biological samples from cell cultures to tissues. The method could also function alongside other modalities like electron microscopy and will provide an exemplar but also further opportunities to integrate and develop new multi-modal techniques and tools for cellular high-resolution imaging in the future.

## Supporting information

All Supplementary

## Acknowledgements

We thank Jeff Irwin (Carl Zeiss Inc) for his dedicated and enthusiastic support with all technical issues and applications relating to the TXM, Damian Reynolds (Carl Zeiss Inc) for technical support, Michael Schwertner (Linkam scientific) for cryo-microscopy expertise and support advice, and Eva Pereiro (Mistral, ALBA) for guidance, encouragement and troubleshooting with issues on soft X-ray tomography. We also thank staff at both Micron, DLS as well as B24 team members and support personnel (past and present) for their help and perseverance. Special thanks to Adam Taylor, Adam Prescott for technical support on a day-to-day basis, Martin Harrow for IT support, Stephen Graham for contributing to our user protocols and Liz Duke for initiating the X-ray tomography project.

M.A.P. was funded via the Wellcome Centre for Human genetics by the Wellcome Trust ISSF grant 105605/Z/14/Z. The Wellcome Trust grant 203141/Z/16/Z funds the Wellcome Centre for Human Genetics. I.M.D. was funded in part by Wellcome Strategic Awards 091911 and 107457 (PI Ilan Davis). P.P-G. acknowledges CROCOVAL ANR-18-CE45-0015 and is part of the national infrastructure “France BioImaging” supported by the ANR PIA1 ANR-10-INBS-04. J.M.G. was supported by Wellcome Investigator Award 200835/Z/16/Z. S.B. was supported by Heisenberg grant: BO 4340/1-1 and SFB1129: Project number 240245660 (Project 14 of SFB 1129) from Deutsche Forschungsgemeinschaft (DFG). M.L.S. was supported by the Brigitte-Schlieben Lange Program from the state of Baden Württemberg, Germany and grant number STA 1536/2-1 from the DFG. DIS is supported by Diamond and MRC grant MR/N00065X/1.

## Author contributions

I.K. collated and evaluated results, processed data and produced the manuscript. M.L.S., S.B., J.M.G. and M.H. designed, performed and analysed reovirus work. M.A.P. and I.M.D. developed cryoSIM design, installed and commissioned of optomechanical components, control algorithms and software (based on original design by John Sedat). M.H., I.M.D., D.I.S. and I.D. conceived this correlative imaging project for the beamline. M.H., I. K. and J.M.G. supervised experiments, collected and processed imaging data on both the cryoSIM and the TXM. P.P-G, M.H. and X.H. did the *in silico* 3D alignment of imaging data. H.W. provided optics support for the X-ray microscopy. C.A.O., T.M.F. and M.C.S provided technical support at beamline B24. All authors contributed to manuscript writing.

## Additional information

Supplementary Information is available for this paper

Correspondence and requests for materials should be addressed to M.H.

## Data availability

Imaging data described in this manuscript will be deposited with the Image Data Resource https://idr.openmicroscopy.org and will be released to the public upon publication.

## Methods

### CryoSIM optical setup

Excitation path: The illumination path consists of dichroics and mirrors to combine 4 linearly vertically polarised illumination beams (405nm, 488nm, and 647nm Omicron Deepstar; 561nm COBOLT Jive^TM^; the polarisation angles are carefully matched using half-wave plates). The combined light is then passed into a telescope section with a pinhole at the focus to act as a spatial filter. This infinity focused beam is delivered to the main beam height (200mm) and passed through a square aperture onto a variable phase delay nematic liquid crystal SLM (SLCoS-LM, Meadowlark, USA) used as a phase grating to produce structured illumination patterns. The light reflected from the SLM is refocused by a lens (all lenses are achromatic doublets) to produce an array of diffraction spots. An aperture at the focus of this lens blocks light from higher order diffraction spots, allowing orders 0 and ±1 through. These pass through a half waveplate and then an LCD based polarisation rotator (PR, Meadowlark, USA) to maintain radially linearly polarised spots (necessary to ensure good structured illumination pattern contrast in the image plane). Another telescope transfers the spot images on to a silver coated mirror, passing through a primary dichroic (Chroma ZT 405-488-561-647-22.5deg, USA). This dichroic transmits the excitation light and reflects the emitted fluorescence. The reflected beam passes through another telescope and reimages the focused diffraction spots in the back focal plane of the objective. A 45° mirror reflects the beam down onto the sample through an 100X air objective (Nikon CFI TU Plan Apo EPI 100X, 0.9NA, 2mm working distance).

Emission path: Fluorescence signal from the sample is collected by the objective and transmitted back from the 45° mirror, through a 1:1 telescope and reflected by the silver-coated mirror upstream then separated from the excitation light by the primary dichroic. The emitted light is then reflected towards the cameras (iXon Ultra 897, Andor) by a broadband dielectric mirror and a final telescope which magnifies the image to optimise the camera pixel size to the optical resolution. There is a secondary dichroic, that splits the light before the detectors. Each detector has a dedicated filter wheel to allow selection of different emission channels. All hardware is controlled by two open source software packages, Cockpit and python-microscope. These packages allow control of complex microscope systems in real time and provide a simple, user-friendly interface.

CryoSIM layout, optics, hardware and software will be reported in detail separately (manuscripts in preparation).

### Virus production and purification

Reovirus T3D strain was produced by infecting suspension of L-cells with a T3D stock originally obtained from B. N. Fields. Virus particles were pre-purified from L-cells by sonication and freon (1,1,2-trichloro-1,2,2-trifluoroethane; Sigma) extraction; virus particles were then purified through ultracentrifugation on Cesium Chloride (CsCl) gradient and stored in virus buffer (150mM NaCl, 10mM MgCl2, and 10mM Tris-HCl, ph 7.5) as previously described^64^.

### Virus labelling

100μl of reovirus particles (from 10^13^ particles/ml stock) were mixed with 0.4μl of Alexa488 or Alexa647 NHS Ester (8mM starting concentration) (Thermo Fisher Scientific) for 1h at room temperature (RT). To remove unbound fluorophores, virus particles were then purified by gel filtration (7K molecular weight cut-off, Invitrogen).

### Cell culture and cell lines

U209 cells (from ATCC), or U209 cells expressing galectin3-mCherry (a kind gift from Harold Wodrich, Bordeaux) were kept in Dulbeccos’ modified Eagle medium (DMEM; Gibco) media containing 10% foetal bovine serum and 1% vol/vol of penicillin and streptomycin (Gibco) at 37°C and 5% CO_2_. Within the U2O9 population presented in this work, only a proportion of the cells expressed endogenously fluorescent Gal3 (designed to be selected under antibiotic control; no antibiotics were used in this case leading to a mixed population). Suspensions of L-cells (from ATCC) for virus production were maintained in Joklik MEM medium (Sigma-Aldrich) supplemented with 1% L-Glutamine (Gibco.), 2% FBS (Gibco), 2% Neonatal calf serum (Gibco) and 1% vol/vol of penicillin and streptomycin (Gibco) at 35°C. BSC-1 cells (obtained from ATCC) were cultured and maintained in DMEM supplemented with 10% fetal bovine serum and 1% penicillin/streptomycin.

### Live-cell imaging of virus infection

U209 wild-type or stably expressing mCherry-Gal3 were transfected with an eGFP-Rab7 expressing plasmid 16h prior to imaging. MRV labelled with Alexa Fluor 647 was added to cells and then imaging was started. Live-cell imaging was performed with an inverted spinningdisk confocal microscope (PerkinElmer) using oil immersion objectives (60x, 1.49 NA, Apo TIRF, Nikon or 100x, 1.4NA, Plan Apo VC, Nikon) and a CMOS camera (Hamamatsu Orca Flash 4). Cells, objectives and microscope stage were kept at 37°C and 5% CO_2_ through the presence of an environment-control chamber. Cells were imaged in 0.5μM stacks 5min apart for 180min.

### Virus infection for X-ray imaging

U209 cells stably expressing mCherry-Gal3 were seeded onto TEM grids (Quantifoil AU G200F1 finder) 16 hours prior to infection. MRV labelled with Alexa-488 was added to cells along with 250nm gold nanoparticle fiducials and grids were frozen in liquid nitrogen-cooled liquid ethane using Leica EM GP2 plunge freezer with a 2s blotting time at 1h intervals. BSC-1 cells were also seeded on TEM grids 16 hours prior to infection. Cells were infected with MRV at an MOI of 100 and 16 hours post-infection grids were frozen in liquid nitrogen-cooled liquid ethane using a Leica EM GP2 plunge freezer with a 2s blotting time.

### Monitoring infection status

Infection prevalence was confirmed via confocal microscopy (presence of green fluorescence virus components intracellularly) before sample vitrification but also with inspection of the same signal once vitrified using a Linkam cryo-stage on a conventional Zeiss microscope (AxioImager) using a 50x objective. The latter allowed us to map grids (using the software LINK supplied with the cryo-stage) and assess their quality with respect to population density, vitrification, presence of fluorophores and grid integrity.

### Cryo-SIM imaging and high-resolution data reconstruction

Vitrified samples on grids were transferred to the cryo-SIM and brightfield imaging was first employed to generate mosaics, where individual cells were evaluated based on cell location (likely to allow data collection on both this instrument and the TXM) and overall state (no obvious grid surface or cell sample disruption). Samples were then imaged in both green and red fluorescence to identify individual cells within the population that both expressed fluorescent Gal-3 and were infected with fluorescent virus. 3D-SIM data was collected on a number of these representative cells (4 mock-infected controls, 8 at 1h PI, 11 at 2h PI, 11 at 3h PI and 5 at 4h PI). Data were reconstructed with SoftWoRX 6.5.2 (GE Healthcare) using real optical transfer functions generated from 3D-SIM images of 175μm single-colour fluorescent beads (PointSpec, Thermo Fisher Scientific) to produce super resolution image stacks. Multi-channel images were aligned with Chromagnon^54^. The raw and reconstructed data were analysed in Fiji^43^ using SIMcheck^44^ to ensure the results were realistic and contained no artefacts.

### Cryo-soft X-ray Tomography and X-ray data reconstruction

X-ray data were collected with an UltraXRM-S/L220c X-ray microscope (Xradia now Carl Zeiss microscopy GmbH) at beamline B24 (DLS) using 500 eV X-rays. This instrument is fitted time with a capillary condenser, a 40nm zone plate objective (25nm for the BSC-1 work) and a 1024B Pixis CCD camera (Princeton instruments). Samples were loaded into the microscope chamber in batches of four and were assessed for structural integrity and alignment potential, inspecting them first with the in-line visible light 20x objective (images recorded on a Retiga 4000R camera; Teledyne) and then with the X-ray microscopy to generate 2D mosaics (visible light gives an overall map of the grid and using X-rays map areas in grid boxes that contained ROIs. The visible light microscopy setup benefits from a variable visible light LED which was used to confirm fluorophore presence and agreement with the fluorescence signal recorded in the cryo-SIM. Tilt series were collected from −65° to +65° at increments of 0.5° on 3 mock-infected cells, 4 cells at 1h PI, 8 cells at 2h PI, 4 cells at 3h PI and 5 cells at 4h PI (all previously imaged at the cryoSIM). All data was aligned and reconstructed automatically to tomograms using the in-house pipeline which employs Batchruntomo^51^ via a number of directives to allow parallel and near real-time processing. Specifically, three protocols for alignment and reconstruction are currently provided via these directives: weighted back projection (WBP), simultaneous iterative reconstruction techniques (SIRT) and automatic patch tracking. Batchruntomo^51^ is part of the IMOD package (version 4.9.2)^46^. Where necessary, multiple adjacent sites were used for data collection for subsequent stitching and FOV expansion.

### Correlative cryo-SXT/ cryo-SIM imaging

Equivalent cryo-SIM and cryo-SXT data sets were overlaid with the multidimensional registration software eC-CLEM^52^. 2D X-ray mosaics were aligned to 2D grid mosaics from the cryo-SIM data collection using the 2D rigid registration mode of eC-CLEM^52^ (coarse alignment). The rotational matrix derived was applied to the 3D X-ray tomograms which were then 3D aligned to the SIM data from that same region using the 3D rigid registration mode of eC-CLEM^52^ (fine alignment). Features used for alignment purposes included both the nanoparticles added on samples just before vitrification as well as grid patterns (grid bars, Quantifoil holes and imperfections) and cellular features that displayed good contrast in both X-ray and light microscopy (fluorescent endosomes in this case).

