## Supplementary material for "Correlative cryo-structured illumination fluorescence microscopy and soft X-ray tomography elucidates reovirus intracellular release pathway": All Supplementary

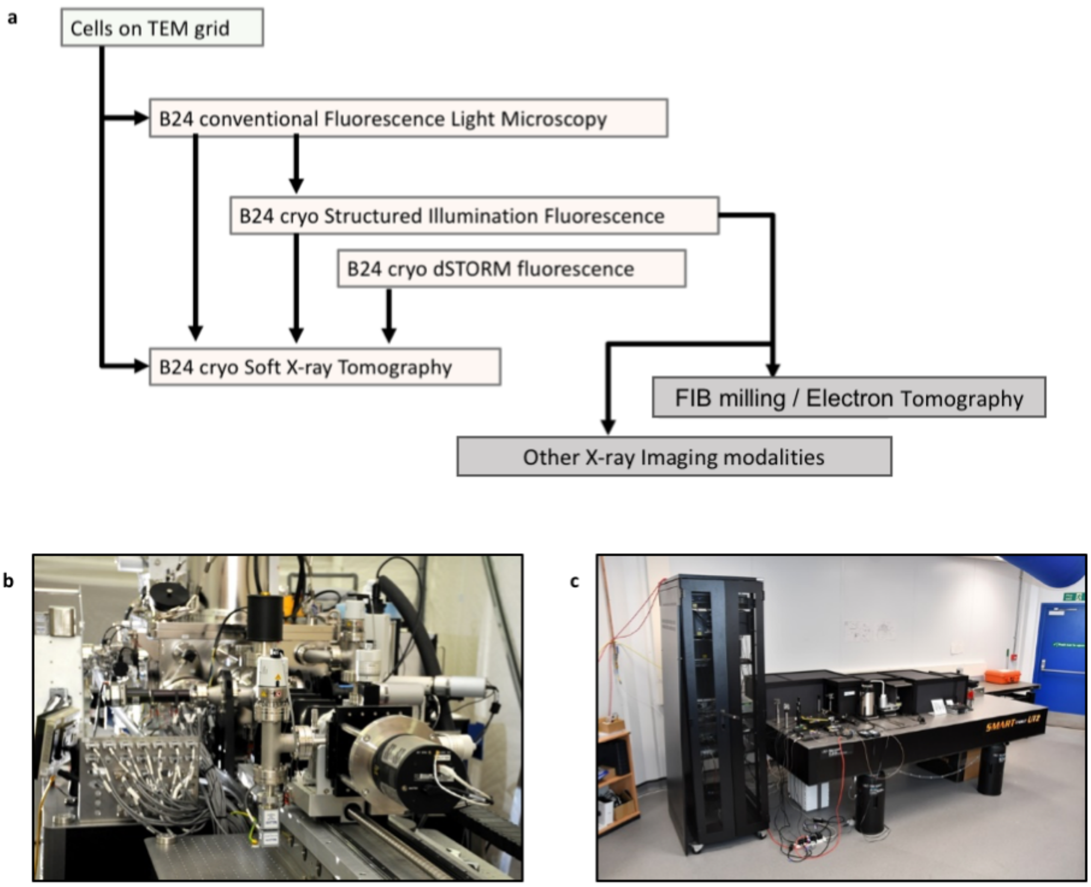


**Supplementary Figure 1**. **Correlative microscopy at beamline B24**
**a**, A schematic of the established workflow through SIM and SXT and including additional functionality available (cryo-dSTORM; currently under commissioning) and potential cross-over to other techniques such as EM and X-ray fluorescence for further correlative imaging. Images of **b**, the TXM and **c**, the cryoSIM on site at the DLS beamline B24 (https://www.diamond.ac.uk/Instruments/Biological-Cryo-Imaging/B24.html).

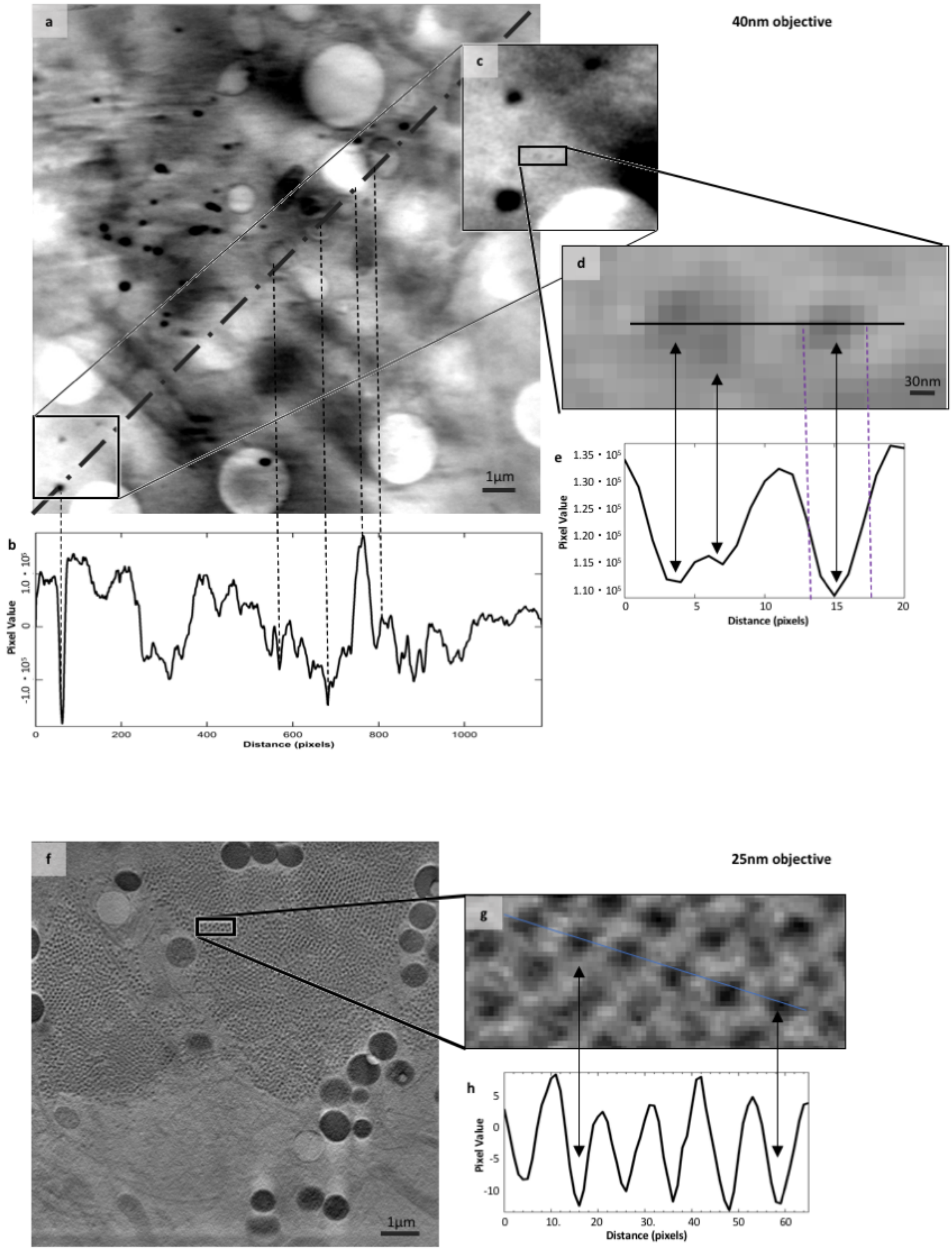


**Supplementary Figure 2. Data contrast in cells using different TXM objectives**

**a**, X-ray projection of a 16x16µm FOV collected at the beamline B24 TXM using the 40nm objective in a U2O9 mammalian cell sample 1h after it has been exposed to infectious reovirus at MOI of 50. The grey arrow points to the cytoplasmic membrane and the zoom in square at the bottom left of the image is located outside the cell. The backing surface (Quantifoil$^{TM}$) can be identified by the regular pattern of circles (needed for blotting prior to vitrification). The diagonal dotted line defines the line used to generate **b**, a line profile of pixel intensity in Fiji[43]. Dips in the profile correspond to features of high X-ray absorbance such as membranes and organelles (corresponding vertical dotted lines have been added as examples). We expect reovirus particles to be present both in and outside the cell but given the high background it is be difficult to unambiguously identify. **c** and **d**, Closeups of marked areas showing contrast that could be attributed to reovirus particles. The associated line profile **e**, clearly contains peaks that are 4-5 pixels apart; at 16nm per pixel that gives an object diameter of approximately 80nm which matches the expected diameter of a reovirus particle. However, the ratio of pixel values is at best 1.1:1.3 even in this 'ideal' area (low background without any overlap with cellular content) making the unambiguous identification of individual virus impossible in this sample. **f**, X-ray projection of a 10x10µm FOV collected at the B24 TXM using the 25nm objective in the cytoplasm of a BSC-1 cell 16h after it has been exposed to infectious reovirus at MOI of 50-100. The observed regular pattern of small dots is indicative of viral factories with each of these dots representing a single viral particle. **g**, A close up of the boxed area in **f**, with a line drawn to generate **h**, a line profile of pixel intensities. Full width half maxima of this plot gives an average diameter of 60nm for these particles which appear to be an array of immature particles in the viral factory having yet to assemble an outer capsid before cell exit.

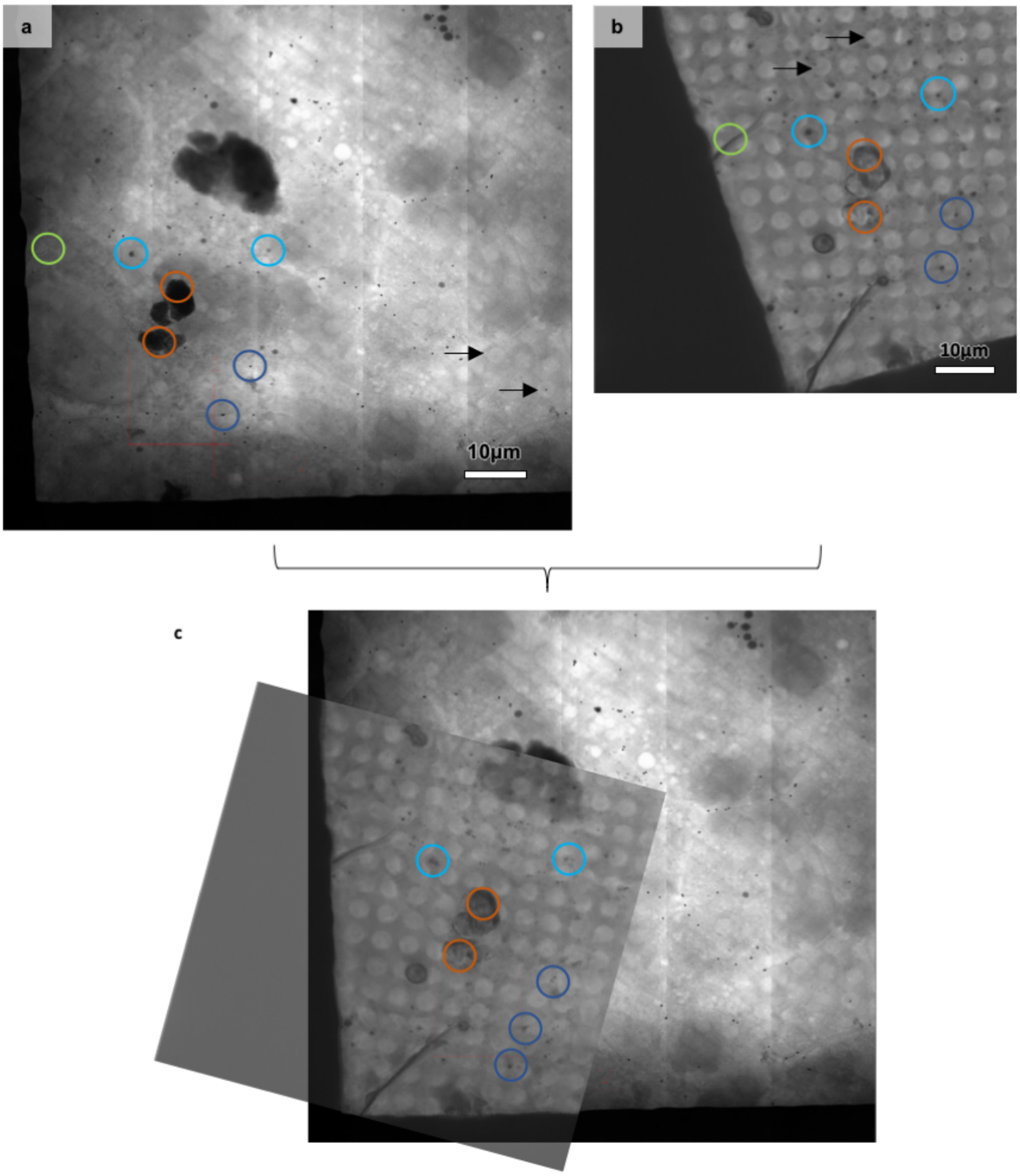


**Supplementary Figure 3. Sample features used as alignment features**

An example of the use of sample and holder features to estimate data associations for alignment purposes. Here, gold nanoparticles (in light blue circles), lipid droplets (in dark blue circles) and surface amorphous ice contaminants (in orange circles) as well as holes in the carbon support film and grid bars can be seen in both **a**, the 2D X-ray mosaic and **b**, the visible light brightfield slice from the cryoSIM and can be used to guide data preliminary overlay shown in **c**.

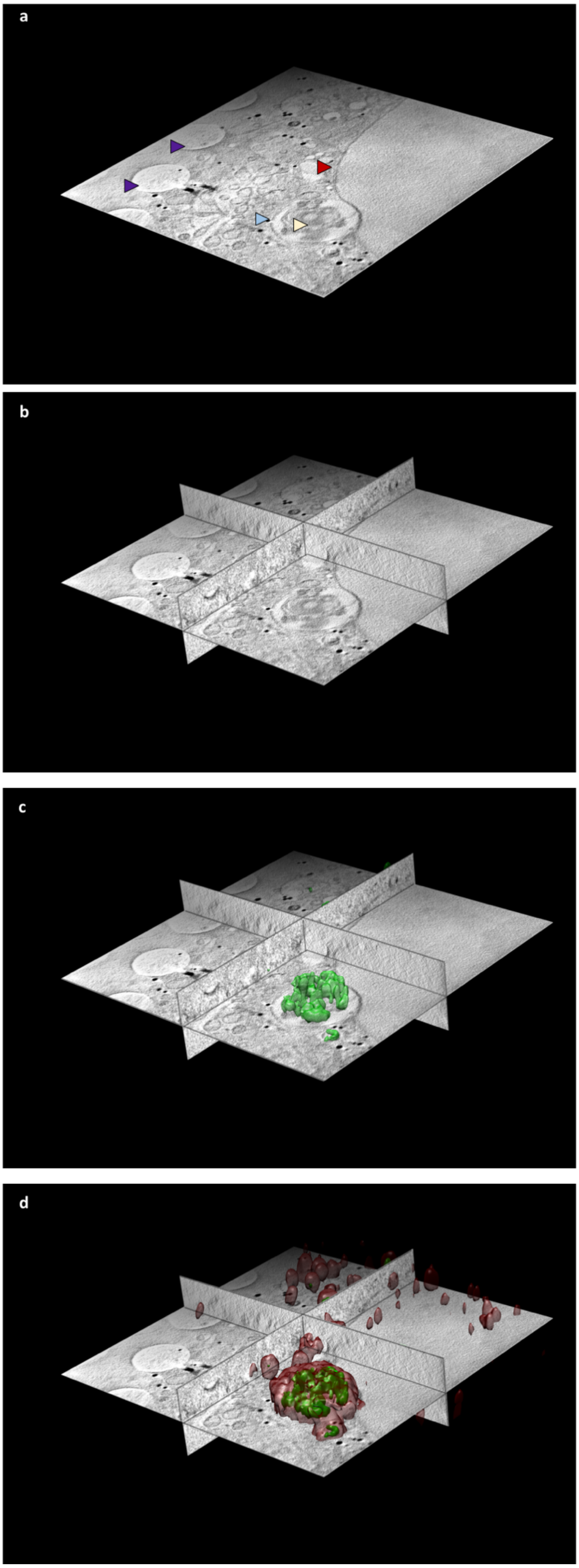


**Supplementary Figure 4. Correlative imaging at beamline B24**

All the data shown here are from a U2O9 cell vitrified 4h PI with high titres of reovirus T3D. Viral components are fluorescently labelled with Alexa488 (green fluorescence) and there is also an endogenously expressed reporter molecule (Galectin-3) which is accumulated in vesicles carrying concentrated reovirus outer capsids and which carries the mCherry fluorophore (red fluorescence). **a**, Z slice from the middle of a soft X-ray absorption tomogram, from beamline B24 using the 40nm objective, of a U2O9 cell with the nucleus on the righthand side of the figure (red arrow; the two leaves of the nuclear membrane are seen in incredible detail), and the cytoplasmic area with a substantial multi-vesicular body/endolysosome (pale blue arrow), containing folded carbon-dense material given the increased X-ray absorption in that structure (pale yellow arrow). In the thinner areas of the distal cytoplasmic region several holes can be seen as part of the support film used for cell attachment (deep purple arrows). **b**, Ortho-slices of **a**, showing a representative portion of the volumetric data. **c**, 3D rendering of the cryoSIM recorded structures displaying green fluorescence within and around the endolysosome and **d**, 3D semi-transparent rendering of all cytoplasmic vesicles that contain red Galectin-3 fluorescence (data also collected on the cryoSIM) and are seen as an indication of viral presence in those vesicles. All images were generated with Chimera[55].

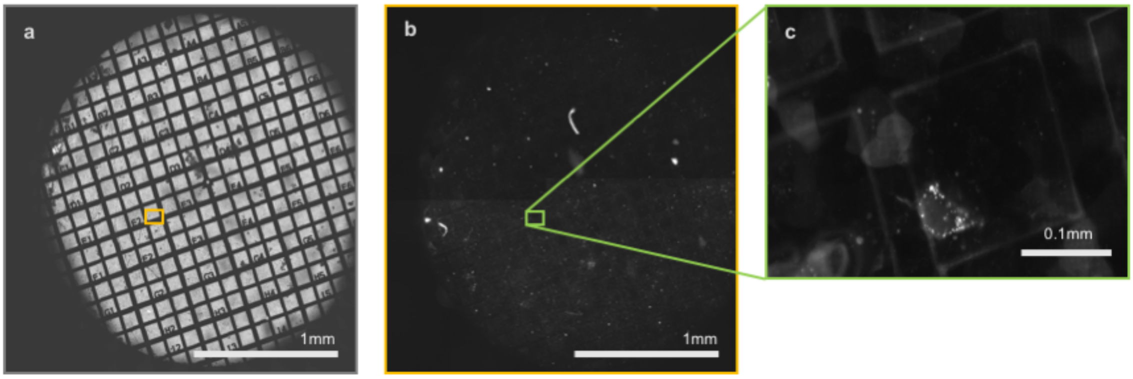


**Supplementary Figure 5**. **Conventional brightfiled and Fluorescent cryo-imaging at B24**
**a**, Brightfield mosaic of a 3mm finder EM grid using white light; **b**, mCherry fluorescence in the same FOV and **c**, expanded view of an area in **b** (box outlined green) with evidence of subcellular structures with red fluorescence (mCherry-Gal3 in this case). Bright white spots are endosomes containing mCherry-Gal3 and conveniently outlining the cell in which they are expressed.

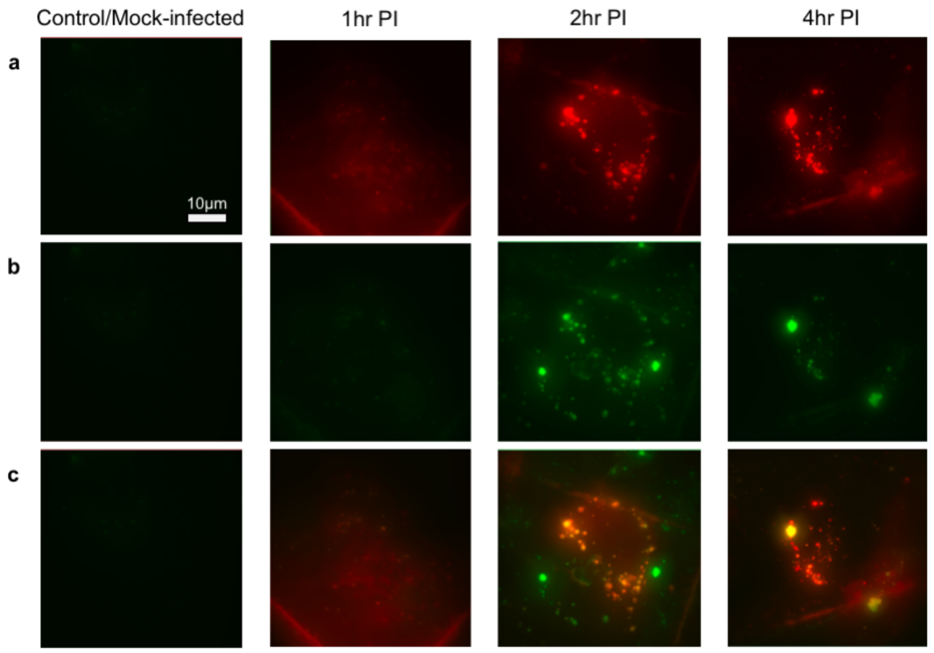


**Supplementary Figure 6**. **Intracellular virus tracking using the cryoSIM**
Representative pseudo-widefield, maximum intensity projections of reconstructed cryoSIM data from reovirus infected cells at 0, 1, 2 and 4h PI with **a**, showing the relative intensity of mCherry (Gal3) at all time points; **b**, Alexa488nm signal (reovirus) and **c**, composite of **a** and **b**. Imaging in each column comes from the same sample/cell and all instances shown here have the same XY spread so the value bar visible in the first panel of row a applies equally to all panels.

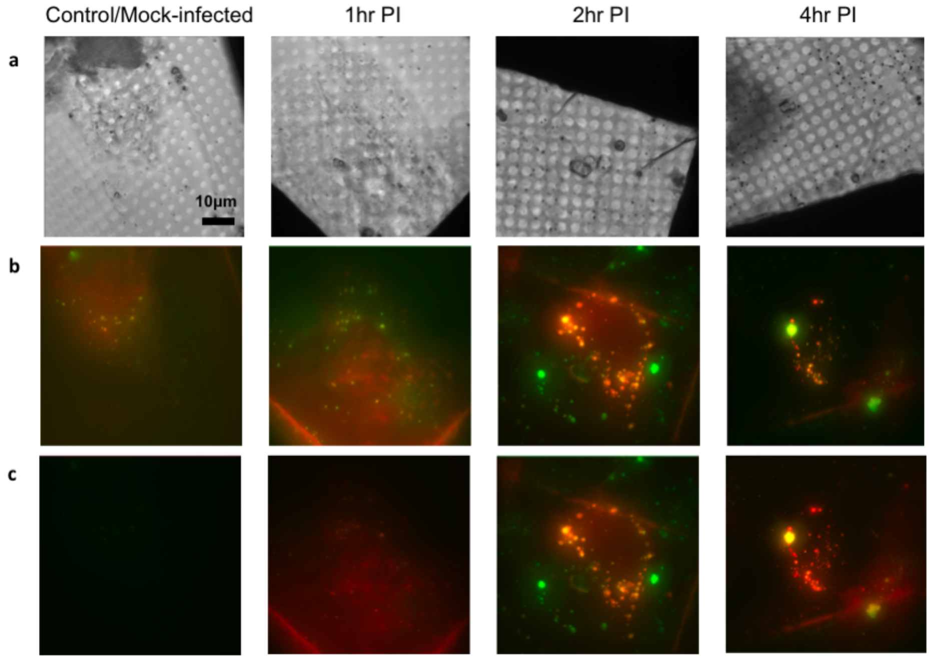


**Supplementary Figure 7. Prevalence of fluorescence localisation after infection**
Gal3 and MRV signal in the cytoplasm of U2O9 cells during early infection stages with **a**, brightfield images of representative cells (Z-axis sum of minimum-intensity projections) at 0, 1, 2 and 4h PI, **b**, maximum-intensity Z-axis projection of reconstructed 3D SIM data at the same time points and **c**, as **b** but all intensity is normalized against the strongest signal recorded at 4h PI to allow us to see the relative increase in signal due to fluorophore localization (Alexa488-MRV in green; mCherry-Gal3 in red). Imaging in each column comes from the same sample/cell and all images are to the same scale.

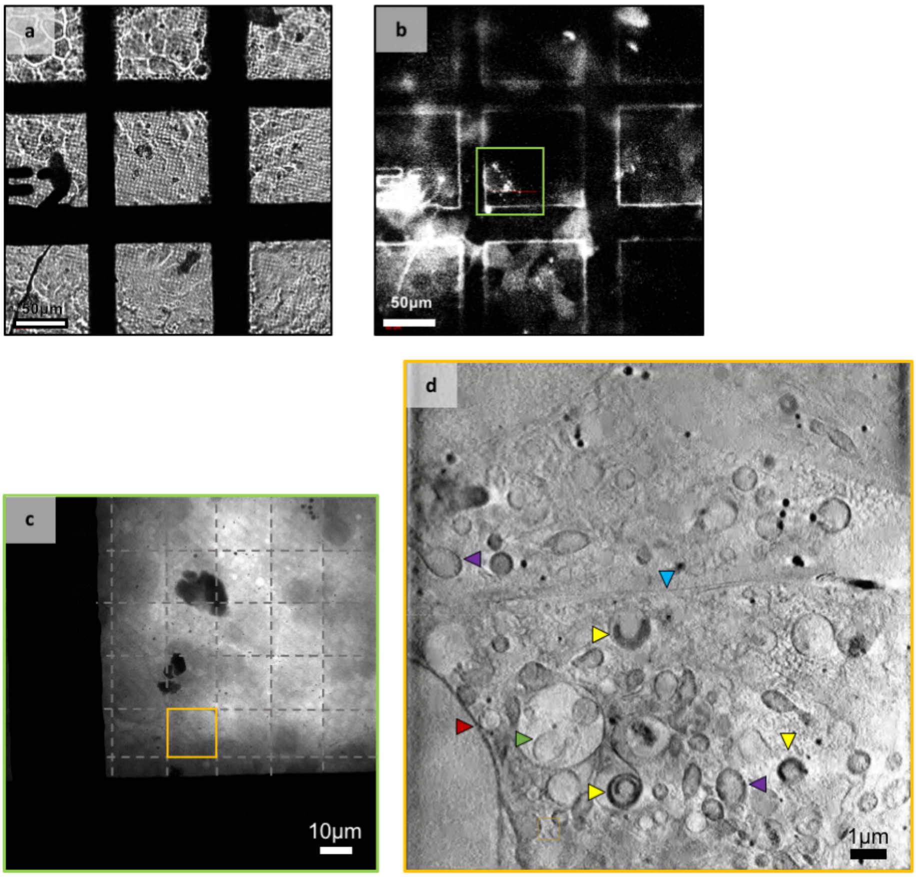


**Supplementary Figure 8. Imaging steps at the B24 TXM**

**a**, Brightfield and **b**, fluorescence in line visible-light imaging of a sample grid supporting a mixed fluorescence cell population. **c**, 2D X-ray mosaic in the ROI denoted in **b** (green outline with individual FOVs delineated by hashed lines); **d**, middle slice from the X-ray tomogram collected in the ROI denoted in **c** (orange outline). Characteristic cellular features are identified by arrows: red for nuclear envelop, purple for mitochondria, green for MVBs, blue for the contact area between two adjacent cells, yellow for MRV-carbon-rich structures within vesicles.

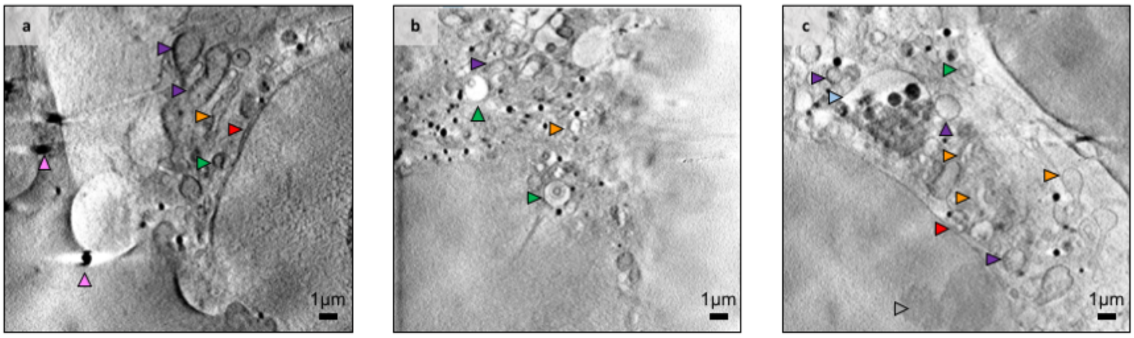


**Supplementary Figure 9. SXT Imaging of control U2O9 cells**

**a-c**, Representative cross-sections from three cells in the control population of U2O9 cells. Cellular structures are identified with arrows: pink for the gold nanoparticle fiducials (250nm), purple for mitochondria, red for the nuclear envelop, orange for the endoplasmic reticulum, green for MVBs, light blue for endo/lysosomes, grey for nucleoli.

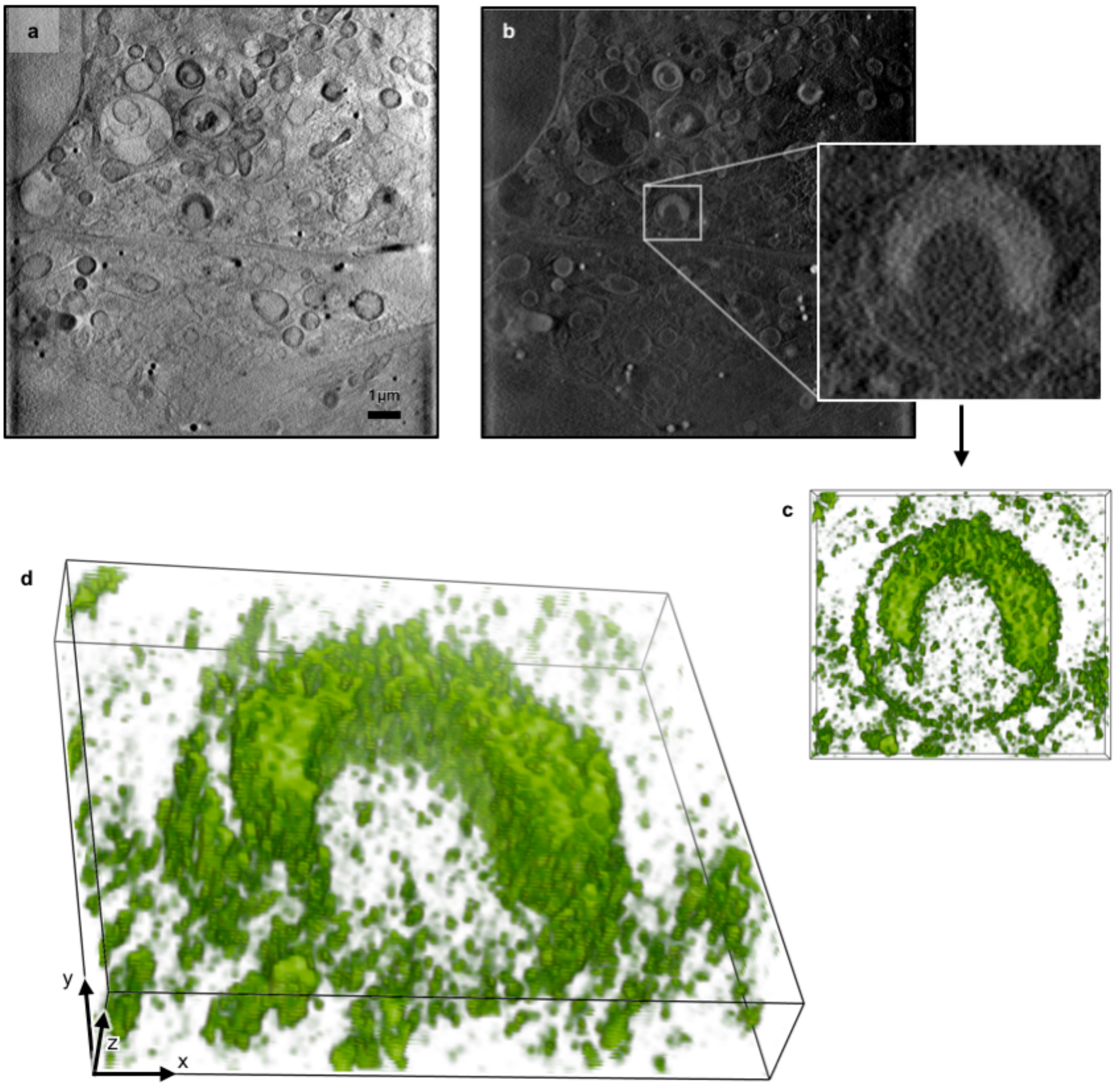


**Supplementary Figure 10. Distinct virus-containing structures in single cytoplasmic vesicles**
**a**, Single Z-axis slice of an ROI X-ray tomogram showing two infected cells at 2h PI (respective nuclei at top and bottom of left-hand side) and **b**, the same with all pixel values inverted (necessary for volume rendering in Chimera[55]) with a sub-area expanded around a distinct semi-circular carbon-dense structure that is reovirus-positive. **c**, Surface rendering of a partial stack of the corresponding density looking down the Z-axis and **d**, the same volume as **c**, tilted to show the 3D structure. All images were generated with Chimera[55].

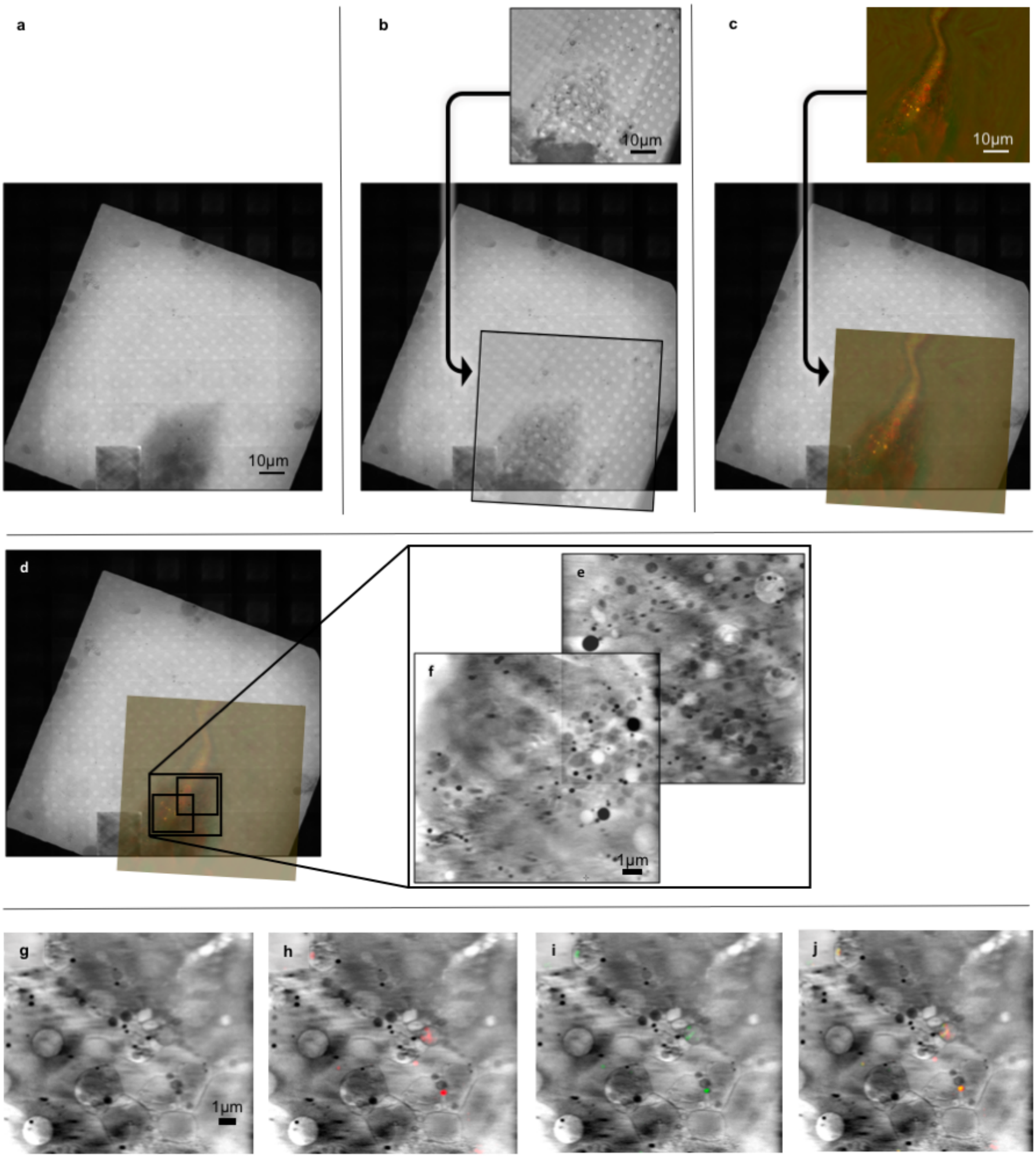


**Supplementary Figure 11. Correlative SIM and SXT on control U2O9 sample (mock-infected)** **a**, 2D X-ray mosaic of a grid square showing a single cell on the south side of the square; **b**, superposition of brightfield slice from the cryoSIM on the same area (pixel interpolation for overlay in eC-CLEM) and **c**, (informed by **b**) superposition of 2D slice from processed SIM data from the same area. The 2D alignment of the above allows **d**, the superposition of all fluorescence data on the X-ray mosaic producing a navigation map for accurate SXT data acquisition. **e** and **f**, Overlapping X-ray tomograms collected in **d,** to allow the 3D stitching of imaging volumes and the effective post data collection increase of the X-ray FOV. **g**, Closeup of representative z slice in the cytoplasmic area with associated **h**, red and **i**, green fluorescence and **j**, their superposition. Both fluorescence signals are perfectly co-localised, weak (relative to those observed later in the infection) and are not associated with any distinct cytoplasmic structures which would imply that they represent the expected background autofluorescence for these cells.

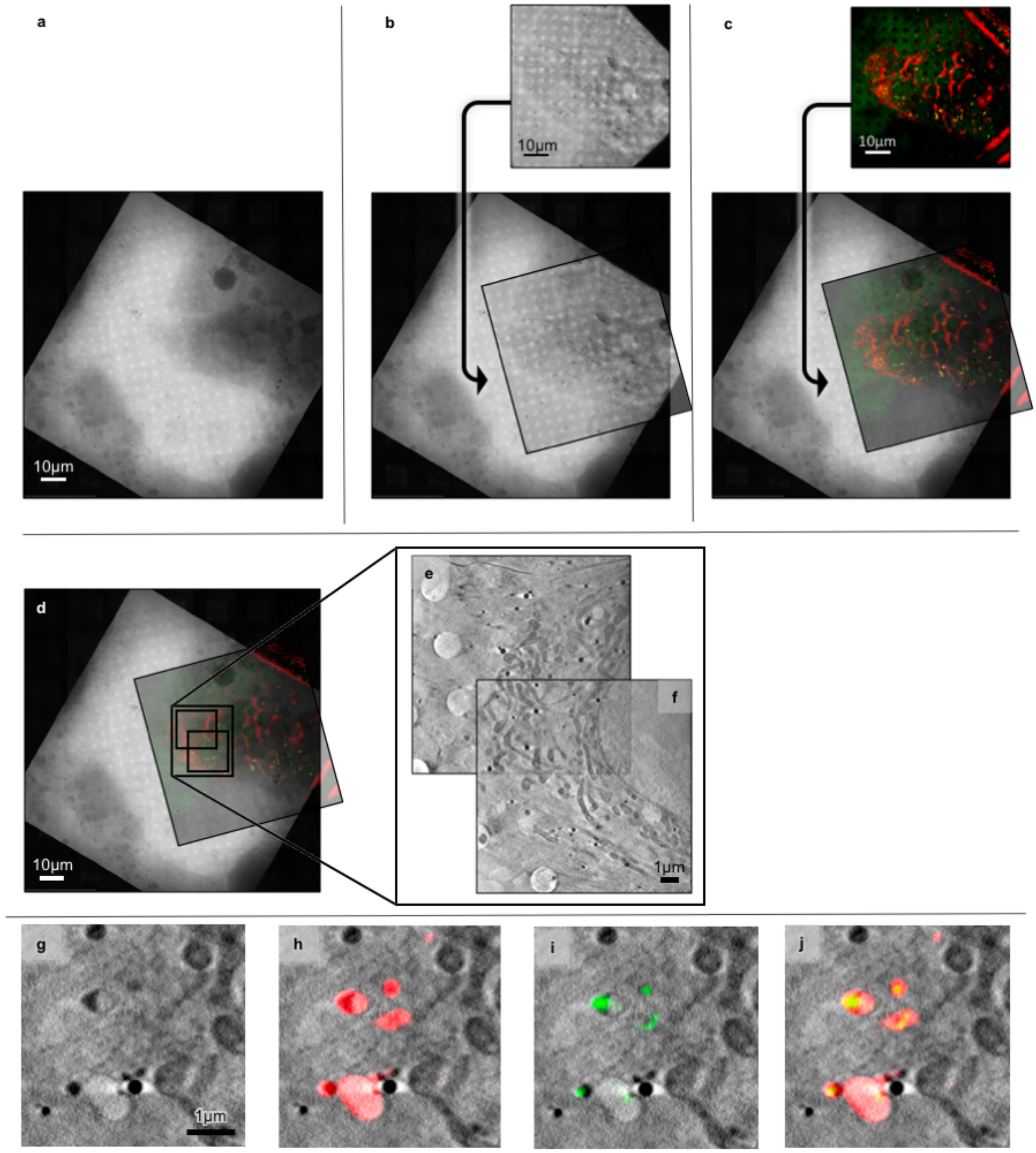


**Supplementary Figure 12. Correlative SIM and SXT on U2O9 cells at 1h PI**

**a-f**, Panels arranged as in Supplementary Figure 11 for a 1h PI sample. **g**, Closeup of cytoplasmic area in the X-ray tomogram shown in **f** with associated **h**, red and **i**, green fluorescence and **j**, their superposition. As shown in **j**, at 1h PI, a number of small but distinct vesicles are evident both Gal3 and MVR positive throughout the cytoplasm.

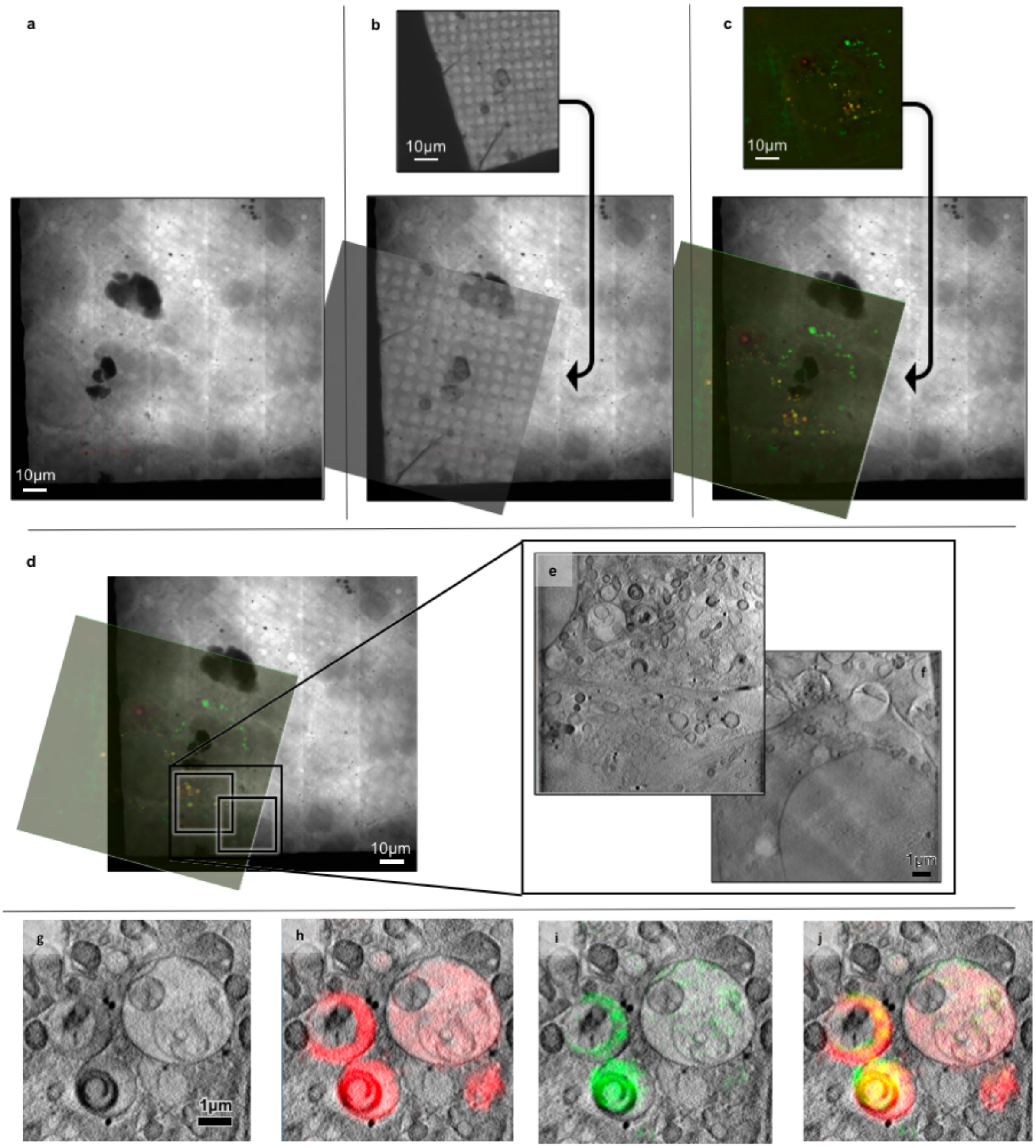


**Supplementary Figure 13. Correlative SIM and SXT on U2O9 cells at 2h PI**

**a-f**, Panels arranged as in Supplementary Figure 11 for a 2h PI sample. **g**, Closeup of cytoplasmic area in the X-ray tomogram shown in **f** with associated **h**, red (Gal3) and **i**, green (MRV) fluorescence and **j**, their superposition. As shown in **j**, at 2h PI, the small vesicles observed by 1h PI have now merged to give rise to MVBs that have distinct compartments infused with Gal3 while others have not been similarly compromised (presumably arising from concatenation of membranes from MRV-free vesicles). Carbon-rich virus-induced horse shoe structures are also seen in single vesicles.

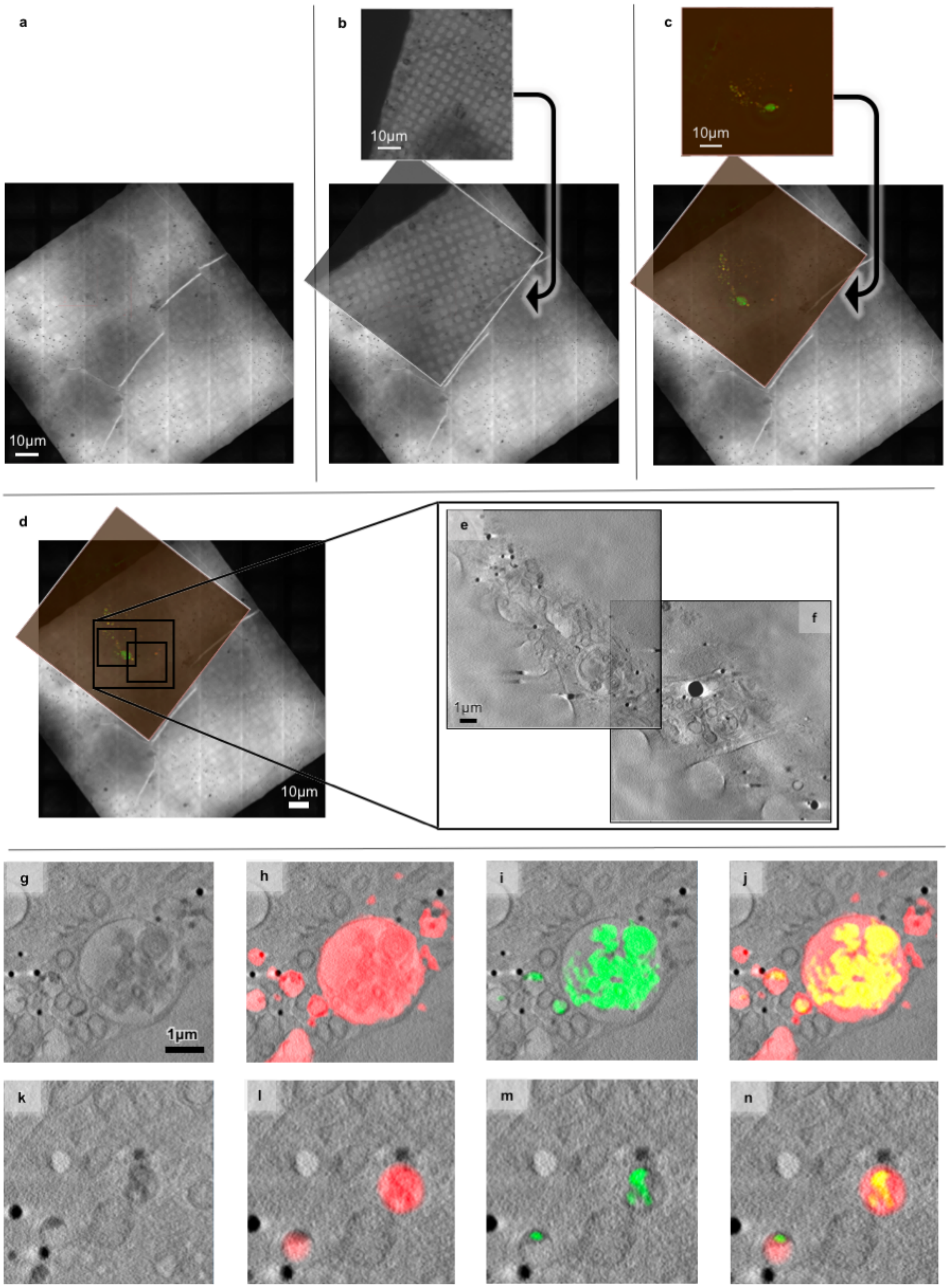


**Supplementary Figure 14. Correlative SIM and SXT on U2O9 cells at 4h PI**
**a-f**, Panels arranged as in Supplementary Figure 11 for a 1h PI sample. **g** and **k**, Closeups of cytoplasmic areas in the X-ray tomograms **e** and **f** respectively with their associated **h** and **l**, red (Gal3) and **i** and **m**, green (MRV) fluorescence and **j** and **n**, their superpositions. Large vesicles such as seen in **g** are typical MVBs after association with lysosomes and further accumulation of cell debris in the process of waste clearance. The folded material inside them is associated with viral capsid components given the colocalization of green fluorescence on the carbon-rich folds. **n**, Another instance of co-localisation exhibits the power of the

techniques used as they allow not only the high-resolution visualisation of the 3D structure of the two organelles in this view but also denote a distinct and highly detailed fluorescence signature on those structures.

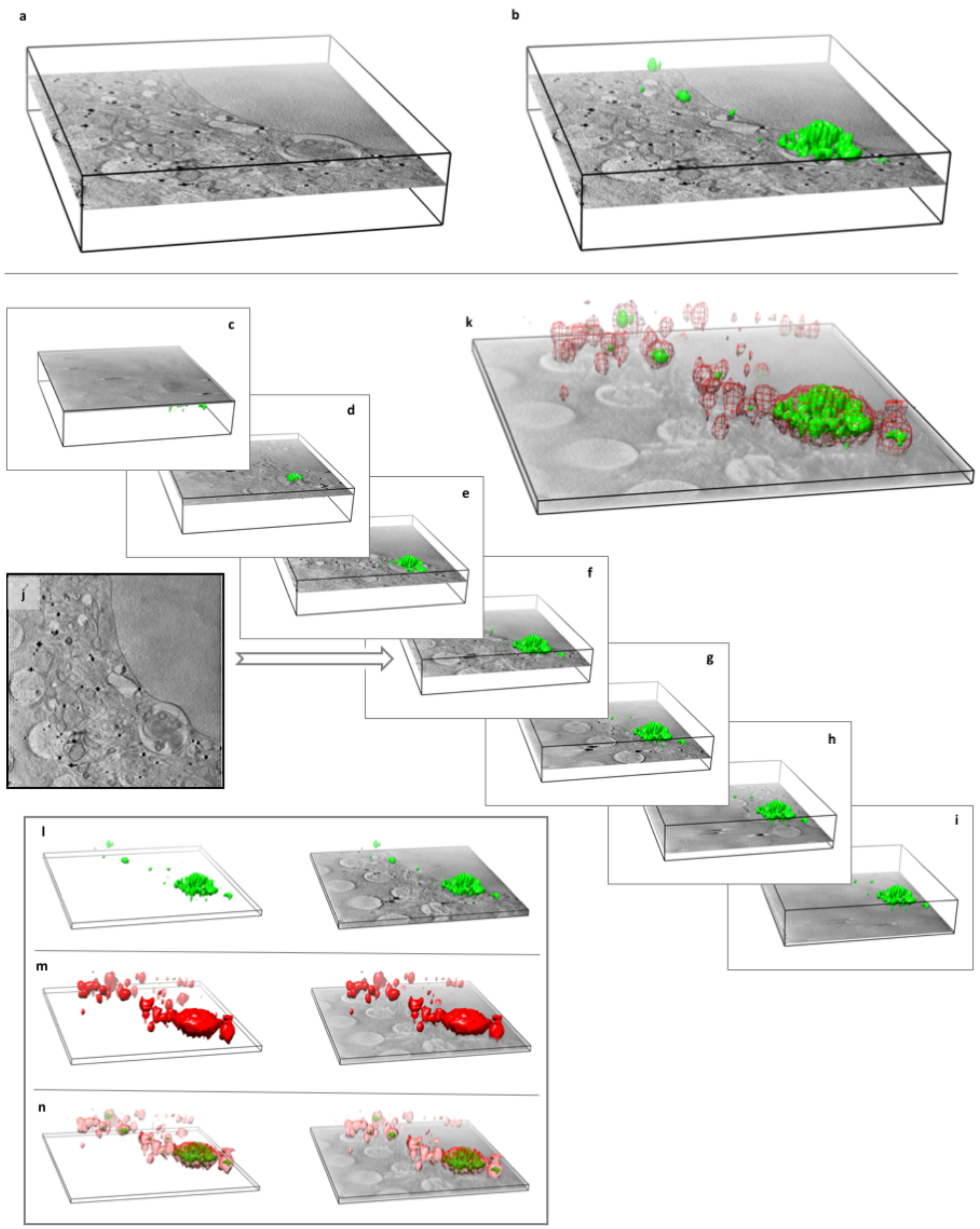


**Supplementary Figure 15. An illustration of the power of the information content of the presented correlative imaging platform**

**a**, Tomographic volume from a U2O9 cells at 4h PI (represented by a delineated box) with a single slice from the corresponding image stack. **b**, Same view as **a**, with the reovirus fluorescence volume (in green). **c-l**, Sequential slicing along the X-ray tomogram Z-axis with the viral volume kept in place. **j**, Single X-ray-only projection of plane represented in **f**. **k**, Reduced X-ray stack with reovirus areas rendered as a green solid volumes and the Gal3 distribution in red mesh representation. **l-n**, Fluorescence volumes in the same tomographic area with only the bottom leaf of this stack to allow an appreciation of the depth perspective of this system of correlative imaginig. All images were generated with Chimera[55].
